# Non-target effects of ten essential oils on the egg parasitoid *Trichogramma evanescens*

**DOI:** 10.1101/2022.01.14.476310

**Authors:** Louise van Oudenhove, Aurélie Cazier, Marine Fillaud, Anne-Violette Lavoir, Hicham Fatnassi, Guy Pérez, Vincent Calcagno

## Abstract

Essential oils (EOs) are increasingly used as biopesticides due to their insecticidal potential. This study addresses their non-target effects on a biological control agent: the egg parasitoid *Trichogramma evanescens*. In particular, we tested whether EOs affected parasitoid fitness either directly, by decreasing pre-imaginal survival, or indirectly, by disrupting parasitoids’ orientation abilities. The effect of Anise, Fennel, Sweet orange, Basil, Coriander, Oregano, Peppermint, Mugwort, Rosemary and Thyme EOs were studied on five strains of *T. evanescens*. Specific experimental setups were developed, and data obtained from image analysis were interpreted with phenomenological models fitted with Bayesian inference. Results highlight the fumigant toxicity of EOs on parasitoid development. Anise, Fennel, Basil, Coriander, Oregano, Peppermint and Thyme EOs are particularly toxic and drastically reduce the emergence rate of *T. evanescens*. Most EOs also affect parasitoid behavior: (i) Basil, Coriander, Oregano, Peppermint, Mugwort and Thyme EOs are highly repellent for naive female parasitoids; (ii) Anise and Fennel EOs can have repellent or attractive effects depending on strains; and (iii) Sweet orange, Oregano and Rosemary EOs have no detectable impact on orientation behavior. This study shows that EOs fumigation have non-target effects on egg parasitoids. This highlights the need to cautiously precise the deployment framework of biopesticides in an agroecological perspective.

## Introduction

Botanical pesticides (BPs) are often presented as an ecofriendly solution for pest management (Regnault-Roger, 1997; Regnault-Roger et al., 2012; Mossa, 2016; Pavela and Benelli, 2016; Isman, 2020). BPs exploit plant allelochemicals for their repellent or toxic effects on many arthropods (Regnault-Roger, 1997; Mossa, 2016). Among the BPs, Essential Oils (EOs), the fraction of volatile fragrant compounds derived from aromatic plants, affect a wide range of insect taxa (Regnault-Roger, 1997; Regnault-Roger et al., 2012) through different neurotoxic effects (Mossa, 2016; Pavela and Benelli, 2016). They can be repellent (e.g. *Foeniculum vulgare* (Bedini et al., 2016), see also Nerio et al. (2010) for a review), antifeedant (e.g. *Ocimum basilicum* (Saroukolai et al., 2014)) or antiovipositant (e.g. *Coriandrum sativum* (Saxena and Basit, 1982)), inhibit digestion, decrease reproduction by ovicidal (e.g. *Origanum vulgare* (Baricevic et al., 2001)) or larvicidal effects (e.g. *Citrus aurantium* (Sanei-Dehkordi et al., 2016); *Mentha piperita* (Morey and Khandagle, 2012); *Thymus vulgaris* (Szczepanik et al., 2012)), or directly decrease adult survival (e.g. *Pimpinella anisum* (Sampson et al., 2005); *Artemisia vulgaris* (Wang et al., 2006); *Rosmarinus officinalis* (Hanane et al., 2018))

BPs, and more specifically EO-based products, are usually considered as low-risk products because they present a low toxicity to non-target vertebrates and show little persistence in the environment (Regnault-Roger et al., 2012; Pavela and Benelli, 2016). However, the non-target effects of these BPs remain poorly known and would need to be accurately assessed (Amichot et al., 2018). Non-target effects of a pest-control strategy include any consequences, different from the desired effect (e.g. death, repulsion, attraction for trapping, etc) on the target pest species. Non-target effects might be helpful (e.g. effects on other pest species) or problematic (e.g. effects on pollinators or natural enemies of the pests), and might affect very different ecological levels from individual organisms to ecosystems (Köhler and Triebskorn, 2013). The effects of BP application on untargeted arthropods for instance, including sub-lethal impacts on beneficial insects, need to be better documented (Regnault-Roger et al., 2012; Haddi et al., 2020; Siviter and Muth, 2020). EOs might actually induce non-target effects against pollinating insects such as bees (Vital et al., 2018), and natural enemies such as parasitoids (Ilboudo, 2009; González et al., 2013; Poorjavad et al., 2014). In an Integrated Pest Management (IPM) context, different strategies can be required for crop protection, and BPs can be deployed together with biocontrol agents such as parasitoids and predators. The efficiency of such programs might thus rely on the innocuousness of these biopesticides for natural enemies.

*Trichogramma* (Hymenoptera: Trichogrammatidae) are tiny egg parasitoids used as biocontrol agents to control many Lepidopteran pests around the world (Consoli et al., 2010; Van Lenteren, 2012). In particular, *T. evanescens* Westwood is commercialized in augmentative biocontrol programs in many crops such as corn, vegetable or sugar-cane (Hassan, 1993). Moreover, *T. evanescens* naturally occurs in many agricultural fields and natural environments in Eurasia (Pintureau, 2009). Non-target effects on *T. evanescens* could thus have both economical and ecological consequences. Regarding EOs, some *Trichogramma* species are sensitive to contact (Parreira et al., 2018a,b, 2019; Alcántara-de la Cruz et al., 2021) or fumigant (Poorjavad et al., 2014) toxicity. Brief immersion of host eggs in diluted EO drastically reduce parasitism by *Trichogramma* wasps (Parreira et al., 2019; Alcántara-de la Cruz et al., 2021). Moreover, exposition of parasitized host eggs to EOs might reduce parasitoid emergence, longevity and fecundity and affect their progeny (Poorjavad et al., 2014; Parreira et al., 2018a,b, 2019; Alcántara-de la Cruz et al., 2021). EO fumigation increases adult mortality and affects mating and oviposition behavior (Poorjavad et al., 2014).

EOs used in IPM programs might affect parasitoids not only directly, because of their toxicity on development, survival and reproduction, but also indirectly, as fragrant products, by disrupting parasitoids’ host searching ability. Indeed, the fitness of egg parasitoids is highly dependent on their ability to locate and recognize their host (Price, 1975). To this purpose, *Trichogramma* depends on many chemical cues both from their hosts and the plants (Consoli et al., 2010; Wajnberg and Colazza, 2013). Foraging females can rely on chemical cues coming directly from the host stage they parasitize, such as compounds present on the surface of the eggs (Frenoy et al., 1992; Renou et al., 1992), or signals from different host stages such as larval frass (Rani et al., 2007), wing scales (Lewis et al., 1975; Ananthakrishnan et al., 1991; Fatouros et al., 2005; Milonas et al., 2009) or sex pheromone (Noldus et al., 1990; Frenoy et al., 1992; Boo and Yang, 1998; Geetha, 2010). *Trichogramma* also exploit chemical signals from the plant emitted either constituvely (Constitutive Volatile Organic Compounds, (Altieri et al., 1982; Romeis et al., 1997; Boo and Yang, 1998; BAI et al., 2011; Wilson and Woods, 2016)) or induced by the presence of hosts such as hosts’ feeding (Herbivore-Induced Plant Volatiles (Peñaflor et al., 2011)) or ovipositing (Oviposition-induced plant volatiles (Fatouros et al., 2012)) behavior.

The objective of this study is to evaluate the non-target effects of 10 essential oils showing insecticidal potential (Anise, Fennel, Sweet orange, Basil, Coriander, Oregano, Peppermint, Mugwort, Rosemary, Thyme) on five strains of the biocontrol agent *T. evanescens*. We focused on fumigation mode because contactless applications seem to exhibit less phytotoxicity on crops, then fumigant formulations of EO-based BP might be favored (Cloyd et al., 2009; Ikbal and Pavela, 2019; Werrie et al., 2020). Two kinds of experiments were conducted to evaluate how EO vapor fumigation affects (i) parasitoid pre-imaginal survival and (ii) adults’ orientation behavior. On the one hand, toxicity was evaluated on parasitoid development by estimating the fumigant toxicity of EOs on pre-imaginal survival. On the other hand, behavioral consequences were studied by testing if vapors of pure EO influenced female parasitoid distribution in a 4-way olfactometer.

## Materials & Methods

### Insects

Five strains of *T. evanescens* were obtained from the Biological Resource Centre Egg Parasitoids Collection “Ep-Coll” (Ris et al., 2018). Strains were chosen among the 20 available living strains of *T. Evanescens* in the BRC Ep-Coll. Strain diversity was maximized by choosing strains from different sampling sites and from different sampled host plants. Strain AF017 had been collected in 2015 on *Olea europaea* in Bergheim (France). Strain AM002 had been collected in 2015 on *Cydonia oblonga* in Estrablin (France). Strain BL065 had been collected in 2016 on *Malus sp*. in Le Change (France). Strain ESP467 had been collected in 2016 on *Phaseolus vulgaris* in Olazti (Spain). Strain MURU222 had been collected in 2016 on *Solanum lycopersicum* in Orthez (France). AF017 and AM002 had been reared on isofemale lines for seven generations. Experiments were carried out in 2018, so parasitoids were reared in the lab for about 30 to 55 generations before experiments.

Parasitoids have been reared in the laboratory on UV-sterilized *Ephestia kuehniella* Zeller (Lepidoptera: Pyralidae) eggs. At each generation, about 500 ± 50 eggs of *E. kuehniella* fixed to a card with diluted glue, were presented to emerging parasitoids for them to parasite. Temperature alterned between 19 ± 1 °C and 25 ± 1 °C according to the experimental schedule, in order to modulate the generation time as needed. Light conditions were L12:D12 photoperiod and humidity was 70 ± 10 % RH.

### Essential oils

We studied the effects of 10 Essential Oils. The EOs were chosen among EOs with insecticidal properties in order to explore Mediterranean plant diversity. Sweet orange (*Citrus x aurantium* var. *dulcis*, Rutaceae) and Peppermint (*Mentha x piperita*, Lamiaceae) EOs belong to the list of natural products acknowledged by the french government for phytopharmaceutical use. Rosemary (*Rosmarinus officinalis*, Lamiciaceae) and Thyme (*Thymus vulgaris*, Lamiaceae) EOs belong to the list of active ingredients eligible for minimum risk pesticide product from the United States Environmental Protection Agency. We chose to explore the Lamiaceae family by including two common aromatic plants: Basil (*Ocimum basilicum*, Lamiaceae) and Oregano (*Origanum vulgare*, Lamiaceae) (Ikbal and Pavela, 2019). We diversified plant families by including Asteraceae: Mugwort (*Artemisia vulgaris*, Asteraceae) (Wang et al., 2006) and three Apiaceae: Coriander (*Coriandrum sativum*, Apiaceae) (Khedr et al., 2013), Green anise (*Pimpinella anisum*, Apiaceae) and Fennel (*Foeniculum vulgare*, Apiaceae) (Dunan et al., 2021). All EOs were obtained from Esperis s.p.a..

The full chemical composition of the different EOs is available in the Supplementary Information S.1. Hierarchical clustering show no particular trend linked to the plant family (Fig S.1). However, Anise and Fennel EOs set apart, probably because they both are mainly composed of anethole, known to exhibit insecticidal activity (Dunan et al., 2021; Sousa et al., 2021). Furthermore, Rosemary and Thyme EOs also distinguish themselves since they are heterogeneous (several major constituents) contrary to the other homogeneous EOs, that are mainly composed of a single majority component (>50%, Table S.1).

### Experimental design

#### Pre-imaginal survival

For each *Trichogramma* strain, the day before the exposure to EOs (D-1), small patches with about 25 ± 7 *E. kuehniella* eggs were exposed during 24h to emerging parasitoids (two cards with five patches each, i.e. 250 host eggs, were placed with about 300 individuals with a sex-ratio around 0.45 ± 0.11). On D-day, each patch was placed individually in a 1cm diameter and 15cm long glass tube (Fig 1). On one extremity, glass tubes were closed with a 4cm wet cotton plug in order to increase relative humidity inside the tube (about 40% RH). On the other extremity, a 4cm dry cotton plug with the treatment that was either nothing (control tubes) or, for each EO treatment, 5*μL*, 10*μL* or 20*μL* of pure Essential Oil. Both cotton plugs remained in place during all the experiment. Full air volume inside the glass tubes was about 5.5*cm*^3^. Tested doses thus correspond to 909, 1818 and 3636 ppm. These values were chosen closed to the estimated LC_01_, LC_10_ and LC_50_ of *Ferula assafoetida* EO on adult female *Trichogramma* wasps (Poorjavad et al., 2014). Each treatment was replicated 10 times, meaning 310 tubes for a single strain. Tubes were placed horizontally, at randomly shuffled locations, on plastic racks, each rack supporting 10 tubes (Fig 1). The 31 racks were placed under a hood with minimal aspiration. There was no artificial light and temperature was about 20.1 ± 0.3 °C.

**Figure 1:**
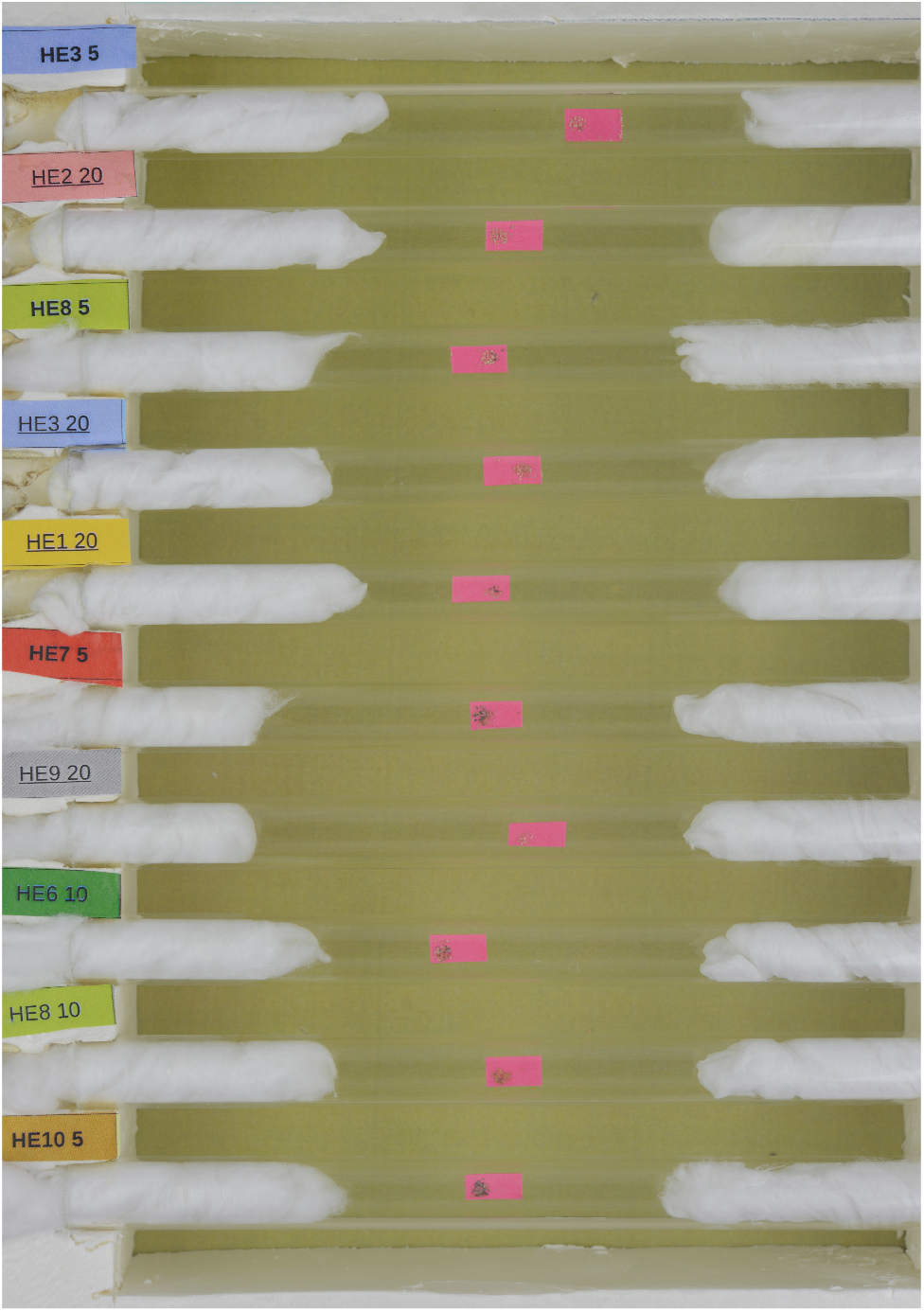
The plastic rack with 10 random glass tubes. Pink cards with egg patches stand inside the glass tubes closed with wet cotton plug on the right side, and Essential Oil treatment (or no additive for control) on the left side. Colored labels identify the Essential Oil and specify the quantity applied (in *μL*).

Five days after having been parasitized, when *Trichogramma* larvae pupate, host eggs turned black (Volkoff et al., 1995). In order to evaluate parasitoid pupation, pictures of the racks were made with a Nickon D800 DSLR on day (D+5). Adults emerged on day (D+14). About one adult emerged per parasitized egg since superparasitism is rarely observed in these conditions, and in most cases, even when several eggs are laid in a single host, a single adult emerge (Corrigan et al., 1995). On day (D+19), in each tube, parasitoids (mostly dead) were counted to evaluate adult emergence.

#### Olfactometry bioassays

Behavioral responses were observed in a four-arm olfactometer (Pettersson, 1970; Vet et al., 1983) inspired, but scaled-up, from the one used by Kaiser (1988). The exposure chamber is a four-pointed star-shape 0.8*cm* thick, sealed on both sides by two 14 × 14*cm*^2^ glass sheets, with fluorocarbon rubber sheet to ensure air tightness. The bottom glass sheet has a hole (5*mm* in diameter) in the center, through which the air flowed out. Full air volume inside the exposure chamber is about 113.1*cm*^3^. The exposure chamber was hold firmly closed with eight pliers (Fig 2). Air circulation inside the exposure chamber has been modelized with Computational Fluid Dynamics to check that the airflow was laminar (see Supplementary Information S.2 for details). This result was confirmed experimentally with smoke tests that show a smooth and regular repartition of the smoke inside the exposure chamber.

**Figure 2:**
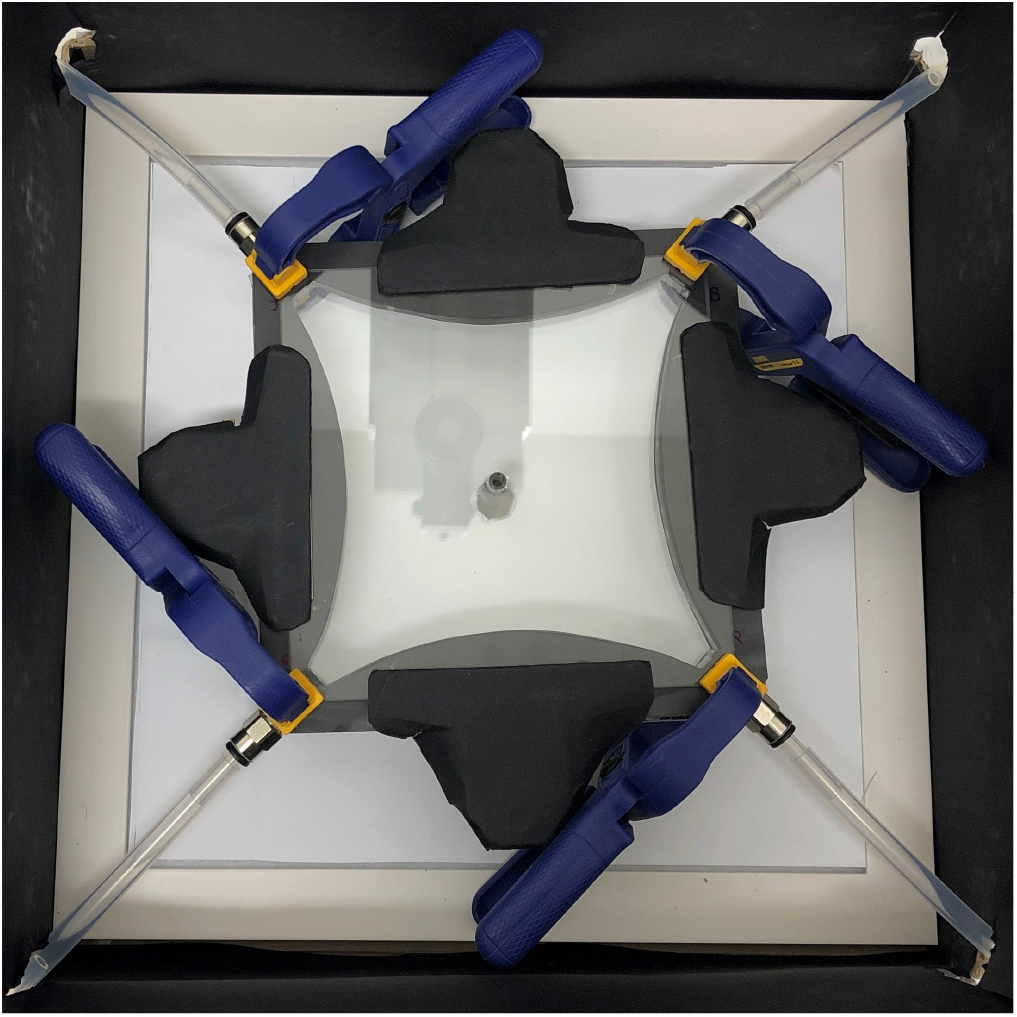
The exposure chamber of the four-arm olfactometer hold closed with pliers was placed on a light pad inside a black box. Air flows from the tubes at the four extremity and evacuate in the central hole.

Air was pushed in the dispositive with an air pump (AquaOxy 2000), filter with Whatman Hepa-vent filter device, and splitted in four arms. Each of the four air flows was set to 1.6*L*.*h*^−1^ with a flowmeter (DK 800 / PV) and sent into 400*mL* distilled water in a 500*mL* Pyrex glass flask. Air from this first flask was sent either into an empty 500*mL* Pyrex glass flask or to a 500*mL* Pyrex glass flask with a VWR filter paper with/without a drop of essential oil. Changing connected flask was made possible with a metal air valve. Both flasks were connected to another metal air valve that commanded the connection with the exposure chamber. In the exposure chamber, air flowed from the four extreme points and created four odour fields that evacuated in the hole in the centre of chamber’s floor. Air from the hole was gently extracted out of the room via an exhaust fan. All connections were fluoropolymere tubes and stainless steel connectors.

Before the beginning of an experiment, the exposure chamber was placed on a led panel (3546 Lumens) and surrounded by 15*cm* high opaque border to avoid any interference from the environment. A DSLR camera (Nikon D810) was fixed 50*cm* above the exposure chamber for image acquisition. About 60 female parasitoids (1 or 2 days old) were brought into contact with the central hole under the exposure chamber. We waited for at least 10 minutes for parasitoids to climb up and enter the exposure chamber. Experiments were stopped (and postponed) when too few individuals climbed up (<10) after 15 minutes. Meanwhile, the Pyrex flasks with VWR filter paper were prepared: two without EO at two opposite corners, and two with the same EO (one with 5*μL* and one with 10*μL*) at the two other corners. EO volumes were chosen closed to concentration potentially used for pest control (e.g. around 20*μL*.*L*^−1^air estimated by Dunan et al. (2021)), but also in order to minimize the risk of toxic effect in the olfactometer (lower than the estimated *C*_01_ for adult parasitoid wasps according to Poorjavad et al. (2014)). For each tested combination strain/EO, four replicates were performed, each time with a different arrangement of the flask locations. After each experiment, all the material was carefully cleaned with 90% ethanol, washed with distilled water and dried. The flasks with VWR filter paper were isolated from air circulation with the metal air valves. When the experiment started, parasitoid movements in the exposure chamber were first recorded during 5 min (hereafter named *control video*). Then, the flasks with VWR filter paper (with or without EOs) were connected to the air circulation and parasitoid movements were recorded for another 15 minutes (hereafter named *treatment video*). The camera settings were ISO 160, F8, with a resolution of 6.7 MP and a framerate of 25 fps.

### Image analysis

#### Pre-imaginal survival

Pictures of host eggs made on day (D+5) were processed with the ImageJ software (Schneider et al., 2012; Schindelin et al., 2012), specifically with the plugin CODICOUNT (Perez et al., 2017). A dedicated macro was written to automatically identify and process each tube in the picture of an entire rack of tubes. The CODICOUNT plugin counted, for each tube, the number of dark and bright pixels in the patch of host eggs, using a segmentation threshold adjusted using manual annotations on a subset of pictures. The two categories of pixels corresponded respectively to black (parasitized) or white (unparasitized) host eggs.

Earlier tests and uses of the method suggested a close linear relationship between the number of pixels and the number of eggs (Burte et al., 2022). The first step was thus to make explicit the link between the number of pixels and the number of eggs. To this end, for each strain, 30 random pictures have been manually counted in order to get both the number of eggs and the number of pixels. Then, for each kind of eggs (black or white), a model was built where the number of black and white eggs (`EggsB`and `EggsW`) was a realisation of a Poisson function whose parameter was a linear function of the number of pixels (`PxB`and `PxW`), such as, for all replicate *k* ∈ [1, 150], 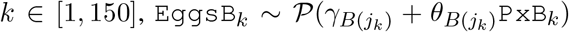 and 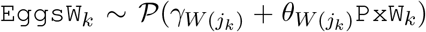. To take into account possible specificity for the strain *j*, parameters *γ* and *θ* were allowed to vary between strains, such as 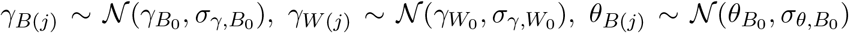 and 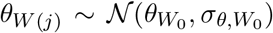 for each strain *j*. All Parameters were estimated with Bayesian inference from non-informative priors (see Supplementary Information S.3.1 for details). From these estimations, both the number of black and white eggs (`EggsB`and `EggsW`) were predicted from the number of pixels for all patches (see Supplementary Information S.3.1 and Burte et al. (2022) for prediction accuracy). The data available after this first step analysis were thus, for all the 310 patches: `NbEggs`, the total number of eggs (`NbEggs`= `EggsB`+`EggsW`) and `NbB`the number of parasitoid pupae `NbB`= `EggsB`).

#### Olfactometry bioassays

Still images were extracted from the video files, using the software FFmpeg: one picture every two minutes for the control videos (three pictures in total) and one picture each three minutes for the treatment videos (six pictures in total). Each picture was manually counted with the assistance of a custom ImageJ macro (Schneider et al., 2012; Schindelin et al., 2012). The location of all parasitoids was marked, as well as the exact location of the chamber (central hole and four extreme points, Fig 3). The ImageJ macro then automatically attributed (x,y) coordinates to each parasitoid, and assigned it to one of 16 portions of the olfactometer chamber, defined with respect to the presence/absence of odors and the shape of the air (see Fig 3). Zones between adjacent fields (i.e. zones with mixed air-flow) were omitted from analyses. The resulting data table contained the number of parasitoids in the whole exposure chamber `NbTotal`(*t*) and in each of the four air fields `Nb`_*z*_(*t*), *z* ∈ {1, 2, 3, 4}, for the different times *t* (*t* ∈ {0, 2, 4} for control experiments and *t* ∈ {0, 3, 6, 9, 12, 15} for treatment experiments).

**Figure 3:**
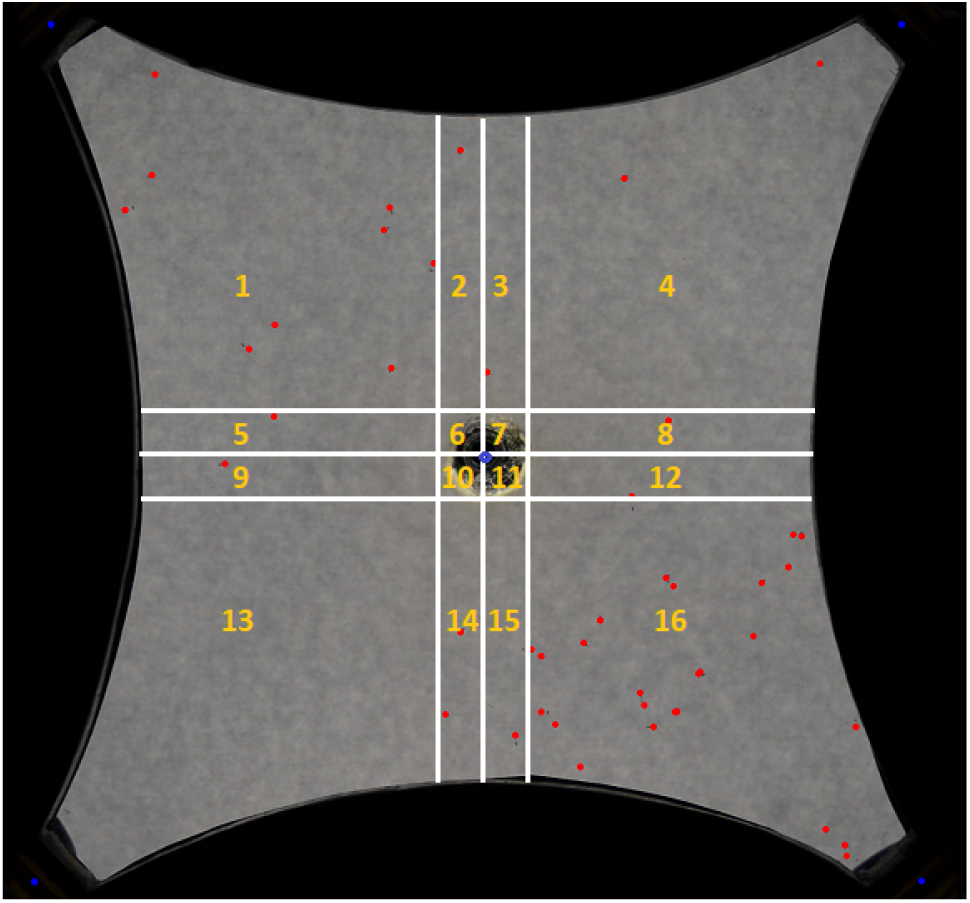
Exposure chamber of the four-arm olfactometer treated by ImageJ. Blue points indicate orientation by identifying the central hole and the extreme points. Red points are manually marked trichograms. 16 zones are automatically defined. To make sure odor fields are homogeneous, adjacent fields (50*pt* horizontal and vertical central strips from the central lines) are not taken into account: only zones 1, 4, 13 and 16 are kept for the analyses (and named zones 1, 2, 3, 4 hereafter). All others zone are gather together and named *y*. This picture is extracted from a treatment video.

### Statistical analysis

Statistical analysis were conducted with the R software (version 3.6.3), using R2jags for Bayesian analyses and ggplot2 for most visualizations (R Core Team, 2020; Su and Yajima, 2015; Wickham, 2016).

#### Pre-imaginal survival

A conceptual model that describes how essential oil affect parasitoid development was built (see Supplementary Information S.3.2 for details). This model includes two steps: (i) parasitism and parasitoid pupation: Are parasitoid eggs laid in the host eggs and do they develop into pupae? and (ii) parasitoid emergence: do parasitoid pupae develop into a living adults that emerge from the host egg?

##### Phase (i) - parasitism and parasitoid pupation

host eggs in a each patch *l* are parasitized with a probability *κ*_*j*_ depending on the strain *j*. Inside the host egg, parasitoid egg survival is assumed to rely on a survival rate that depends on the treatment `C`_*l*_, the volume of EO to which patch *l* was exposed, such as, for five days of exposition to EO, parasitoid early survival probability is 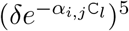. *δ* is natural survival rate. Parameter *α*_*i,j*_ represents daily sensibility to EO *i* for the eggs of strain *j*. Overall, the number of parasitoid eggs turning black is thus 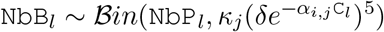.

##### Phase (ii) - parasitoid emergence

adult emergence depends on parasitoid pupal survival probability 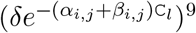. Since adults emerged at (D+14), parasitoid pupae are exposed 14−5 = 9 days to EOs. The number of emerging adults is thus 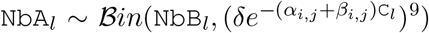. *α*_*i,j*_ + *β*_*i,j*_ is daily sensibility to EO *i* for the pupae of strain *j*. Parameter *β*_*i,j*_ represents the differences between first instars (eggs and larvae) and pupae sensibility: if pupae are more sensitive than larvae, *β*_*i,j*_ > 0, if they are more resistant, *β*_*i,j*_ < 0, and if both pupae and larvae are equally sensitive, *β*_*i,j*_ = 0.

In order to test if the difference of sensibility between the two phases could be explained by a cumulative effect (accumulation of EOs as time passes), a different model structure was tested. In this model, we assumed that daily mortalities were no more independent: instead of representing the survival probability to *n* days of exposition as Survival = (daily survival)^*n*^, we modeled 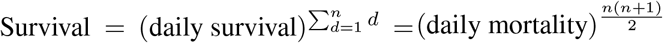 (Fig 4). This alternative model was chosen among other possible functions because the quadratic term allows to represent a form of acceleration. In this model, the effect of EOs was represented by parameter *γ*_*i,j*_ (equivalent of *α*_*i,j*_ in the constant model). In this model, the number of parasitoid eggs turning black becomes 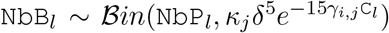 and the number of emerging adults 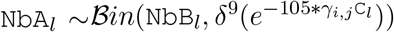. Then, a mixed structure was tested with a model with both an independent daily survival and a cumulative effect. In this model, the number of parasitoid eggs turning black becomes 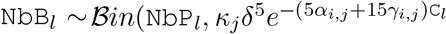 and the number of emerging adults 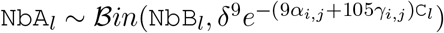.

**Figure 4:**
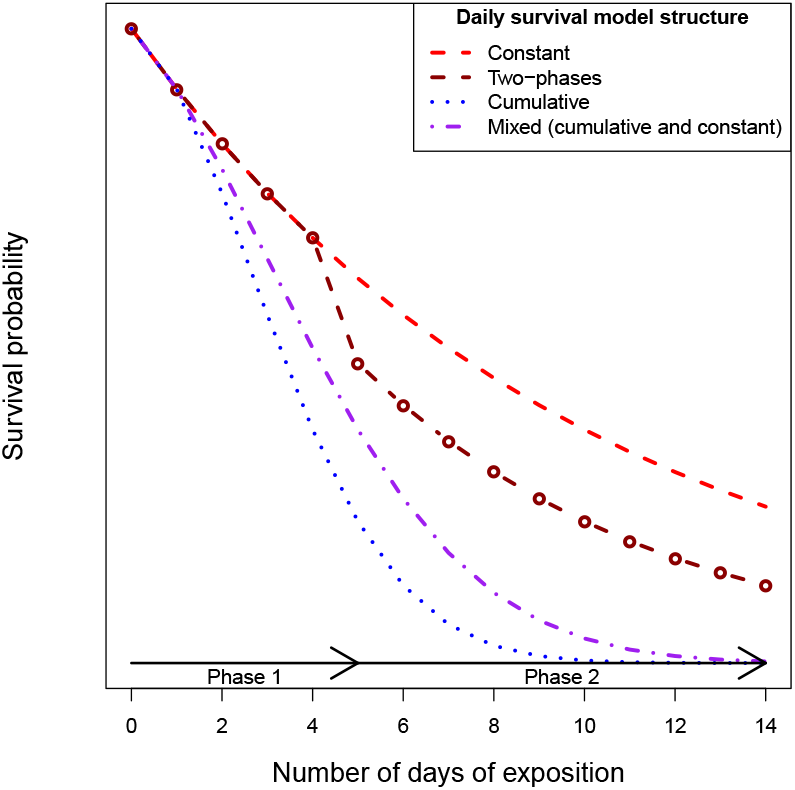
Survival probability according to the number of days of exposition to EOs (*d*) for different model structures. Light red dashed line is independent and constant daily survival (*e*^−0.1*d*^). Dark red dashed line represents independent survival with two phases (*e*^−0.1*d*^, ∀*d* ≤ 5 and *e*^−0.15*d*^, ∀*d* > 5). Blue dotted line is cumulative effect 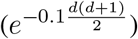. Purple dashed-dotted line is the mixed structure 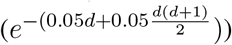.

To estimate the parameters of these models, we performed Bayesian inference (details are provided in the Supplementary Information S.3.2). We tested different possibilities for both parameters *α*_*i,j*_ and *β*_*i,j*_ to study the influence of the EO and the specifity of the strain (Table 2). Parameter *α* was either a sum of EO and strain effects with interaction between both effects (*α*_*i,j*_) or without interaction (*α*1_*i*_ + *α*2_*j*_), or only dependent on EO (*α*1_*i*_). Parameter *β* was either null, or EO dependent (*β*1_*i*_), or strain dependent (*β*2_*j*_), or dependent of both effects either with interaction (*β*_*i,j*_) or without (*β*1_*i*_ + *β*2_*j*_). The best model was chosen by minimizing the Deviance Information Criterion (DIC), a Bayesian measure of fit adequacy, penalized by model complexity (Spiegelhalter et al., 2002; Plummer, 2009).

The effect of EOs were compared by using LD_50_. Global LD_50_ was defined as the dose of EO at which 50% of the parasitized eggs do not transform into adult parasitoids. It was calculated from parameter estimations such as 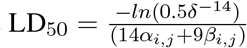. First phase 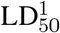 was defined as the dose of EO at which 50% of the parasitized eggs do not develop into pupae (black eggs). It was calculated from parameter estimations such as 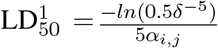. Second phase 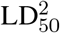 was defined as the dose of EO at which 50% of the parasitoid pupae do not emerge as adult parasitoids. It was calculated from parameter estimations such as 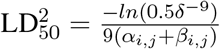.

#### Olfactometry bioassays

We assumed that the presence of EO might only affect the probability for a parasitoid to stay in a given zone of the exposure chamber. In order to estimate this effect, we built a model that represents the parasitoids’ distribution over the four zones throughout an experiment. The parameters of this model were determined with Bayesian inference (details are provided in the Supplementary Information S.4). We first analyzed control experiments to determine parasitoid movements in the absence of EO. Then, parameter estimates from this analysis of control experiments were used as priors in the analysis of subsequent treatment experiments. The model can be divided in four phases: (i) initial distribution of the individuals across the zones, (ii) determination of the portion of individuals that stay in their current zone, (iii) determination of the flow of individuals coming from other zones, and (iv) update of the individuals’ distribution across the zones.

##### Phase (i)

At the beginning of an experiment (either a control or a treatment), parasitoids are randomly distributed in the exposure chamber. The exposure chamber is virtually separated between the four zones with air fields named *z* (*z* ∈ {1, 2, 3, 4}) and all the strips between fields gather together and called *y*. With *μ* being the probability to be in *y*, the number of individuals at time *t* = 0 in zone *z* = 1 is 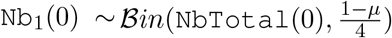. Then, for each *z* ∈ {2, 3, 4}, 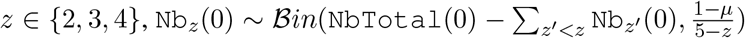. The number of individuals in *y* is thus 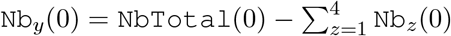.

##### Phase (ii)

In all experiments, a given proportion of parasitoids are assumed to stay in their zone during one minute. This proportion is modelled with an inverse logit function in order to stay in a [0, 1] interval. This proportion is squared for control experiments and cubed for treatment experiments, to make them comparable (since pictures are separated by respectively two and three minutes). In a control experiment, a proportion 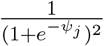 of individuals (from strain *j*) stays two minutes in a given zone. The probability to stay in a zone *z* at time *t* is thus 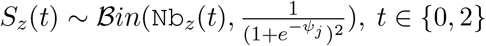, *t* ∈ {0, 2}. In a treatment experiment, the two zones without odor (namely zone 3 and 4) are similar to the control test. The only difference is that the pictures being taken each 3 minutes, the portion of individuals staying in the odorless zone becomes 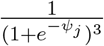. For *z* ∈ {3, 4}, the probability to stay in the zone *z* at time *t* is thus 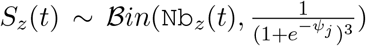, *t* ∈ {0, 3, 6, 9, 12}. The other two zones contain respectively 10*μL* (zone 1) and 5*μL* (zone 2) of essential oil. The probability to stay in these two zones depends on the effect of essential oil *i* on the parasitoids from strains *j* such as 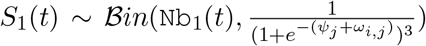, and 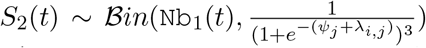, with *t* ∈ {0, 3, 6, 9, 12}. Parameter *ω*_*i,j*_ and *λ*_*i,j*_ respectively represent the effect of 10 and 5 mL of essential oil *i* on strain *j*.

##### Phase (iii)

In both control and treatment experiments, we assume that the parasitoids have enough time (either two or three minutes) to move freely in the exposure chamber. The individuals leaving a given zone are thus randomly distributed between the other zones. For control experiments, the flow of individuals arriving from a zone *z* at time *t* (with *t* ∈ {2, 4}), is 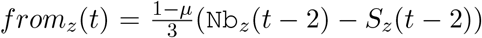. For treatment experiments, the flow of individuals arriving from a zone *z* at time *t* is 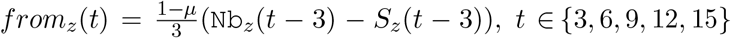, *t* ∈ {3, 6, 9, 12, 15}. In both control and treatment experiments, the flow of individuals arriving from the excluded space *y* is 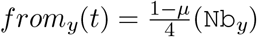, with *t* ∈ {2, 4} in control experiments and *t* ∈ {3, 6, 9, 12, 15} in treatment experiments.

##### Phase (iv)

The resulting number of individuals in a zone is obtained as the realization of a Poisson function depending on the sum of the number of staying individuals in this zone and the flows of individuals from all the other zones. In control experiments, the resulting number of individuals in a zone *z* (*z* ∈ {1, 2, 3, 4}) at time *t* (*t* ∈ {2, 4}), is `Nb`_*z*_(*t*) ∼ 𝒫(*S*_*z*_(*t* − 2) + ∑_*z*′ ≠*z*_ *from*_*z*_′ (*t*) + *from*_*y*_(*t*)). In treatment experiments, the resulting number of individuals in a zone *z* (*z* ∈ {1, 2, 3, 4}) at time *t* (*t* ∈ {3, 6, 9, 12, 15}), is `Nb`_*z*_(*t*) ∼ 𝒫(*S*_*z*_(*t* − 3) + ∑_*z*′ ≠*z*_ *from*_*z*_′ (*t*) + *from*_*y*_(*t*)). In both control and treatment experiments, the updated number of individuals in space *y* is 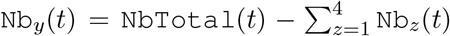. Phases (ii)-(iv) are then repeated until reaching the final time (*t* = 4 for control experiments and *t* = 15 for treatment experiments).

We first tested different kind of parameter *ψ*_*j*_ to analyze data from control experiments with Bayesian inference. *ψ*_*j*_ was either a constant (*ψ*), or variable with strain *j* (*ψ*_*j*_), or specific for each experiments either around a constant value (*ψ*_*k*_) or strain-specific values (*ψ*_*k*(*j*)_) (Table 3). We also checked that no bias existed due to the location by modeling a parameter dependent on the zone of the exposition chamber (*ψ*_*z*_). The pertinence of the whole model was tested by fitting a model *ψ*_−∞_ where no individuals stayed in the different zone and parasitoids were randomly distributed each time step. We relied on the DIC to identify the best control model. From this best control model, we extracted estimated values for parameters *μ* and *ψ*_*j*_, and used them as priors to analyze data from treatment experiments with Bayesian inference (treatment models in Table 4). We tested different parameters *λ*_*i,j*_ and *ω*_*i,j*_. *ω* was either equal to *λ*_*i,j*_ (no dose effect), or to 2*λ*_*i,j*_ (linear dose effect) or independent (non-linear dose effect). Parameter *λ* was either null, or dependent on essential oil without variation between strains (*λ*_*i*_), or dependent on essential oil with fixed variation between strains (*λ*_*i,σ*(*j*)_), or dependent on essential oil with a variance between strains dependent on essential oil 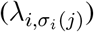. The best model was determined by minimizing the DIC. Details on Bayesian inference are available in Supplementary Information S.4.

## Results

### Pre-imaginal survival

The flexibility of the Bayesian analysis allows to choose between different model structures (Table 1). The structure that best represents the variability of the experimental data considered an independent daily survival probability that separates the toxicity on parasitoid pupation (before egg darkening occurs; *α*) from the toxicity on parasitoid emergence (after the egg has darkened; *α* + *β*). The difference of sensibility between the two phases (represented with parameter *β*) fitted more the data than the accumulation effect as it was modelized (Δ*DIC*_*model*.3,*model*.1_ = 1181 in Table 1). Moreover, the toxicity on both parasitoid pupation and emergence were strain and EOs specific. Indeed, according to the DIC, the best model takes into account *α*_*i,j*_ and *β*_*i,j*_ that both depend on each combination EO x strain (model 1 in Table 2).

**Table 1:** Pre-imaginal survival models according to their daily survival structures (Fig 4). (#) is the number of estimated parameters. Tested models are ranked according to increasing DIC.

| Model | Daily survival model structure | # | DIC |
| --- | --- | --- | --- |
| **1** | **Two-phases** | **106** | **12154** |
| 2 | Mixed (constant and cumulative) | 106 | 13086 |
| 3 | Cumulative | 56 | 13335 |
| 4 | Constant | 56 | 15520 |

**Table 2:**
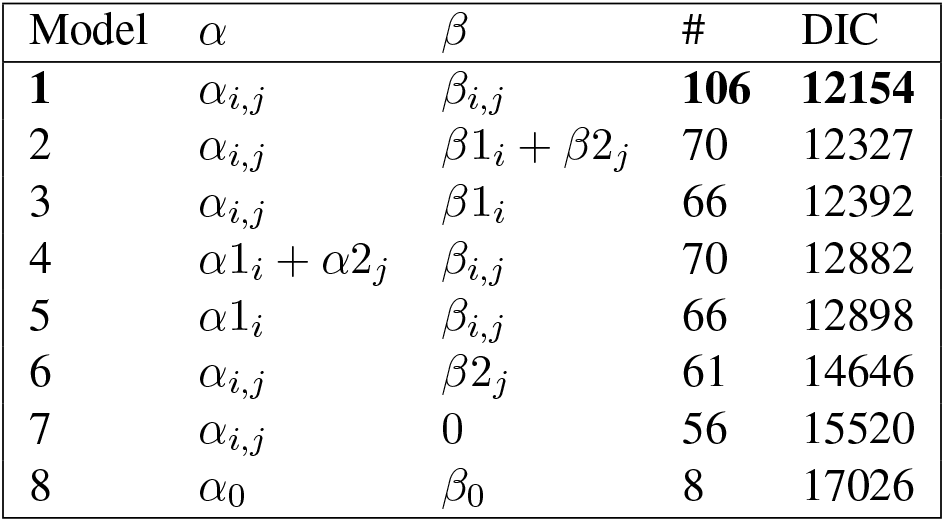
Pre-imaginal survival models according to their structures regarding parameters *α* (toxicity on egg and larval development) and *β* (difference between pupae and first instars sensibility). (#) is the number of estimated parameters. Tested models are ranked according to increasing DIC. *i* and *j* stand respectively for a given essential oil and *Trichogramma* strain.

A large majority of EOs were found to be highly toxic to *T. evanescens* (Fig 5). Indeed, the predicted LD_50_, the dose at which half the parasitoid eggs do not develop into adults is less than 2*μL* for most Eos (Fig 5). Anise, Fennel, Basil, Coriander, Oregano (Fig 6.a), Peppermint and Thyme EOs are extremely toxic for parasitoid pre-imaginal survival since the predicted LD_50_ is less than 2*μL* for all tested strains. Mugwort (Fig 6.b) EO is slightly less toxical with predicted LD_50_ varying between 2*μL* and 6*μL* according to the strain. Rosemary (Fig 6.c) and Sweet orange EOs seem less toxical since their LD_50_ are respectively 10±3*μL*, 6±2*μL* (mean and standard deviation of predicted LD_50_ between the five strains).

**Figure 5:**
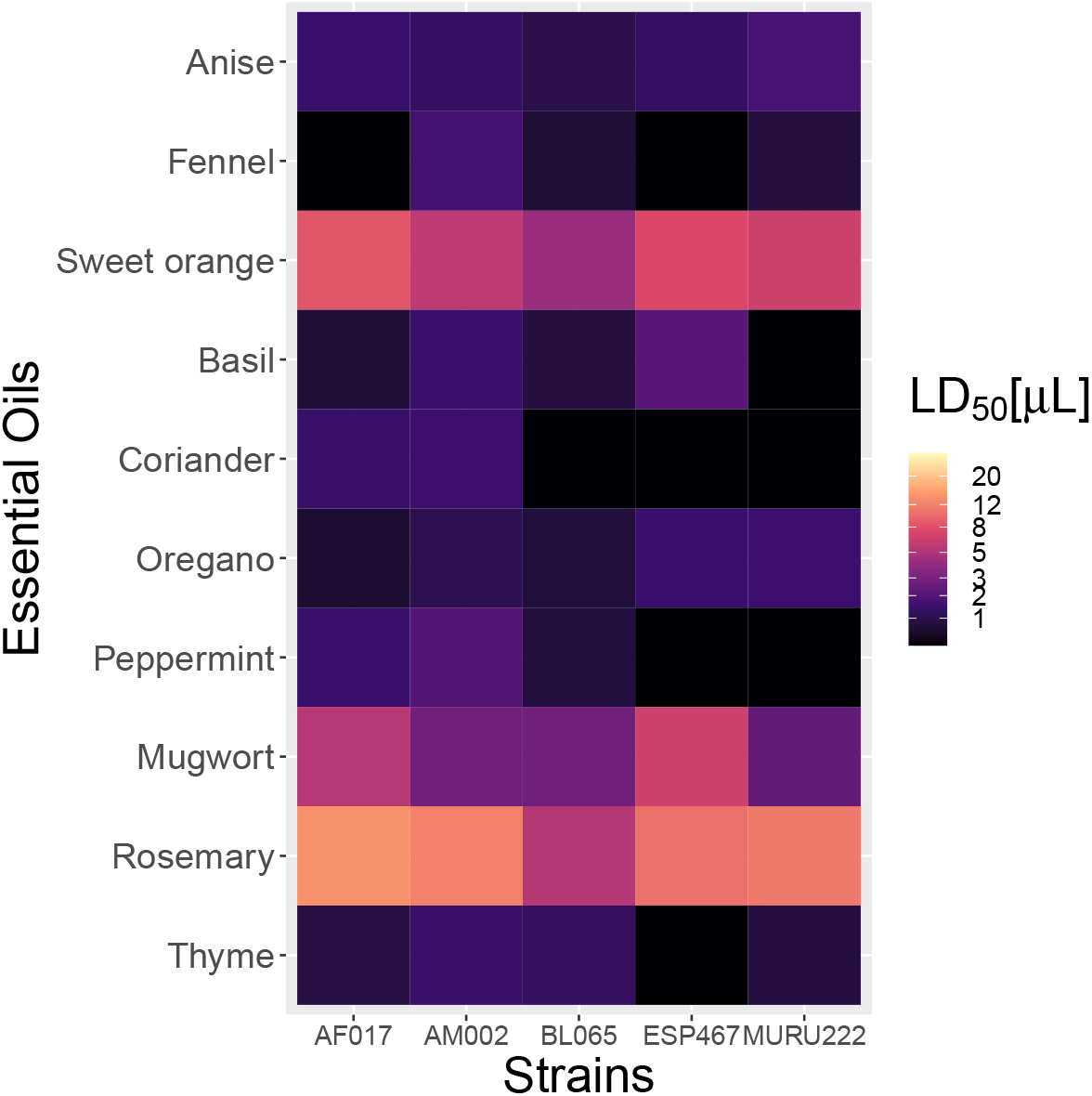
Predicted LD_50_ according to the different strains (columns) and essential oils (rows). Global LD_50_ is the dose of EO at which 50% of the parasitized eggs do not emerge as adult parasitoids. Dark squares represent low LD_50_, meaning highly toxicity of a given essential oil for a given strain.

**Figure 6:**
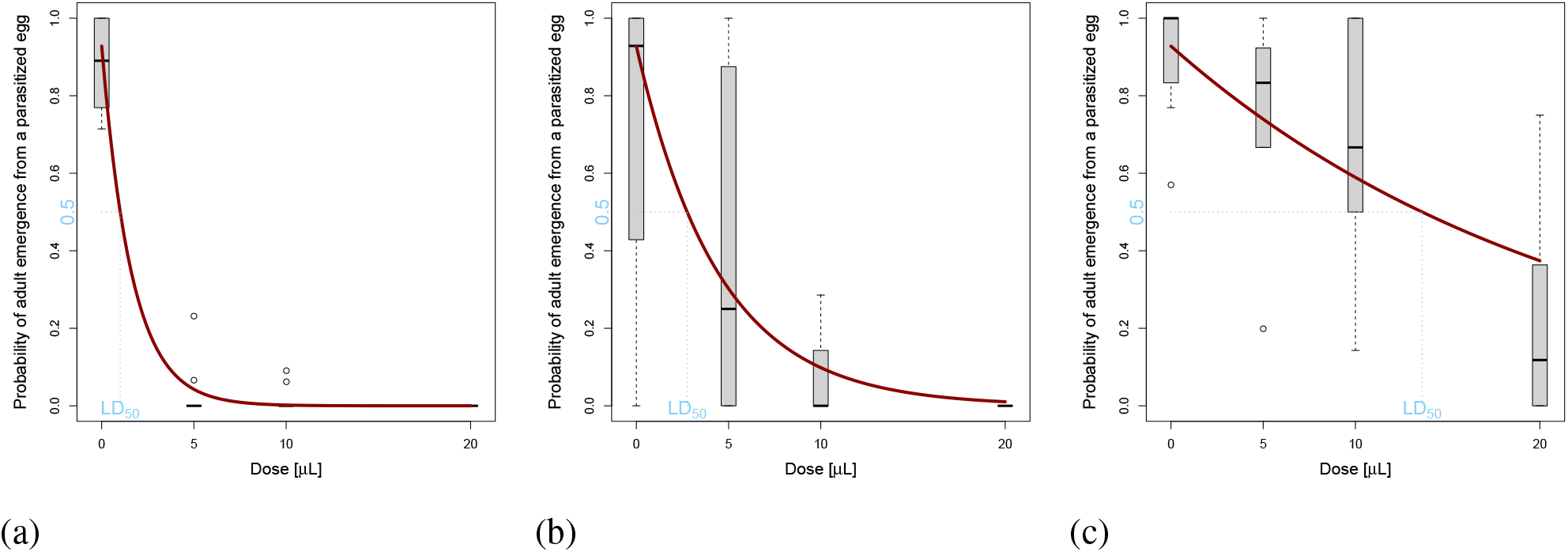
Observed (grey boxplot) and predicted (red line) effect of EOs on the pre-imaginal survival. (a) Oregano essential oil and AM002 strain; (b) Mugwort essential oil and BL065 strain; (c) Rosemary essential oil and AF017 strain. Observations (grey bloxplot) are built by dividing the number of emerged adult parasitoids by the number of parasitized eggs (recomposed by multiplying the total number of eggs by the strain-specific parasitism rate estimates). The survival probability (red line) is estimated from model 1 in Table 2. Dashed light blue lines are the LD_50_ projections.

Toxicity in the first phase of development is variable according to both EOs and strains (Fig 7.a). The interaction between essential oil and strain effects has to be taken into account (ΔDIC_*model*.4,*model*.1_ = 728 in Table 2). For most EOs, the effect on parasitoid survival until pupation is rather low since the predicted 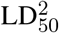 are higher than 25 *μ*L for 42% of the tested combinations (Fig 7.a). However, some EOs appear toxic for this first phase of development: Basil and Peppermint EOs particularly affect egg an larval survival since their 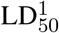 are lower than 25 *μ*L for all tested strains (Fig 7.a). Moreover, strains AF017 and AM002 seem slightly less sensitive than other strains to essential oils in this first phase of development (Fig 7.a).

**Figure 7:**
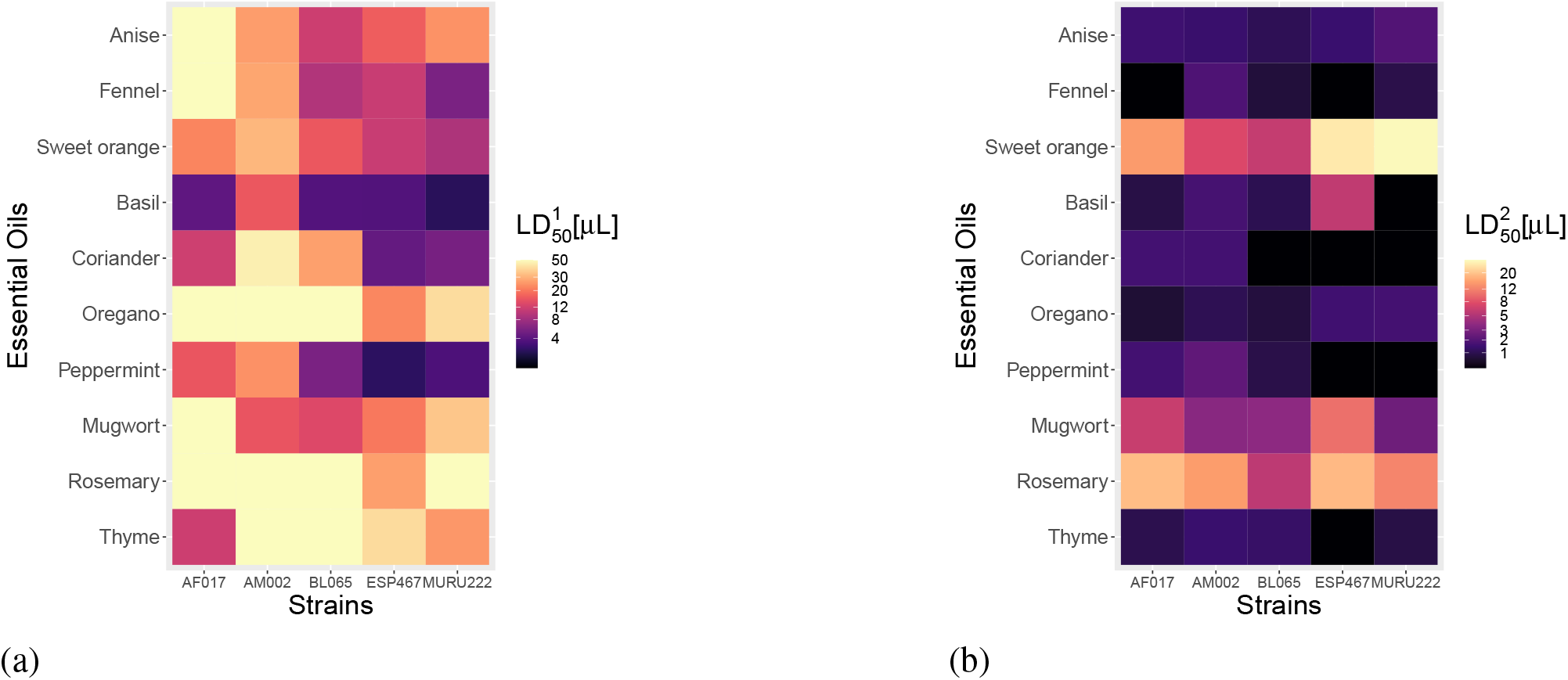
Predicted first phase 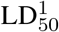 (a) and second phase 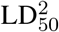 (b) according to both the strains (columns) and the essential oils (rows). (a) first phase 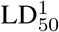 is the dose of EO at which 50% of the parasitized eggs do not transform into pupae. (b) second phase 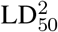 is the dose of EO at which 50% of the pupae do not emerge as adult parasitoids. Lighter squares represent higher values of 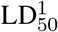 or 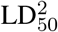, and stand for lower toxicity of a given essential oil for a given strain.

EOs were more toxic on parasitoid development in the second (pupal) phase than in the first (larval) phase (Fig 7). For almost all combinations, exposition to EOs affected more pupae survival than larval survival: for all EOs (apart from Sweet orange EOs for ESP467 and MURU222 strains and Basil for ESP467 strain), the predicted 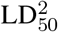 was lower than the predicted 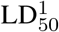 (Fig 7). Beyond the fact that the pupal phase lasts 9 days while the egg-larval phase lasts 5 days, the daily survival was not constant between the two phases (Table 1). The difference of sensitivity between the parasitoid stages was dependent on EOs and strains (ΔDIC_*model*.2,*model*.1_ = 173 in Table 2). In most cases, parasitoid pupae were more sensitive to EOs than larvae since parameter *β*_*i,j*_ > 0 for 94% of all the combinations. Exposition to EOs during this second phase is highly toxic for parasitoids, especially for Coriander, Fennel, Thyme, Peppermint, Oregano, Anise, Basil EOs since their predicted 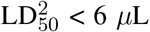 *μ*L for all strains. Rosemary and Sweet Orange EOs seem less toxic for this phase since their predicted 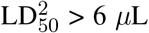 *μ*L for all strains. The pattern observed for this second phase seems to drive the global toxicity pattern on pre-imaginal survival (Fig 5 and Fig 7.b).

#### Olfactometry bioassays

About half of the female parasitoids in the inoculation tube climbed up into the olfactometer chamber (mean = 30 individuals and standard deviation = 10). In control experiments, the portion of individuals that stay in a given zone of the olfactometer chamber did not depend neither on the strain, nor on the experiment, nor on the orientation of the experimental setup (replicate number), as desired (Table 3). Without any EO stimulus injected into the olfactometer chamber, the probability for a parasitoid to stay in a given zone for one minute was estimated around 0.85 (95% posterior credible interval *CI*_95_ = [0.82, 0.87]).

**Table 3:** Control olfactory experiment models according to their structures regarding parameters *ψ* determining the probability for an individual to stay in a given zone for 1 min. (#) is the number of estimated parameters. Models are ranked according to increasing DIC. *j, z* and *k* respectively stand for a given strain, zone or experiment.

| Model | $\psi$ | # | DIC |
| --- | --- | --- | --- |
| **1** | $\psi$ | **2** | **11169** |
| 2 | $\psi_z$ | 5 | 11171 |
| 3 | $\psi_k$ | 3 | 11174 |
| 4 | $\psi_j$ | 6 | 11176 |
| 5 | $\psi_{k(j)}$ | 7 | 11185 |
| 6 | $\psi_{-\infty}$ | 1 | 12885 |

In treatment experiments, the distribution of individuals was strongly impacted by the presence of EOs (ΔDIC_*model*.6,*model*.4_ = 528 in Table 4) but did not depend on the dose of EO (ΔDIC_*model*.3,*model*.1_ = 51 and ΔDIC_*model*.5,*model*.1_ = 160 in Table 4). The effect of EOs was variable across the different strains (ΔDIC_*model*.4,*model*.2_ = 72 in Table 4). Moreover, the inter-strain variance was also dependent on essential oils (ΔDIC_*model*.2,*model*.1_ = 3 in Table 4). The effect of EOs estimated for each strain might be highly variable (Fig 8).

**Table 4:** Treatment olfactory experiment models according to their structures regarding parameters *λ* (odor effect with 5*μL* of essential oil) and *ω* (odor effect with 10*μL* of essential oil). (#) is the number of estimated parameters. Models are ranked according to increasing DIC. *i* and *j* stand respectively for a given essential oil and strain.

| Model | $\lambda$ | $\omega$ | # | DIC |
| --- | --- | --- | --- | --- |
| **1** | $\lambda_{i,\sigma_i(j)}$ | $\lambda_{i,\sigma_i(j)}$ | **22** | **23192** |
| 2 | $\lambda_{i,\sigma(j)}$ | $\lambda_{i,\sigma(j)}$ | 13 | 23195 |
| 3 | $\lambda_{i,\sigma_\lambda(j)}$ | $\omega_{i,\sigma_\omega(j)}$ | 42 | 23243 |
| 4 | $\lambda_i$ | $\lambda_i$ | 12 | 23267 |
| 5 | $\lambda_{i,\sigma_i(j)}$ | $2 \cdot \lambda_{i,\sigma_i(j)}$ | 22 | 23352 |
| 6 | 0 | 0 | 2 | 23795 |

**Figure 8:**
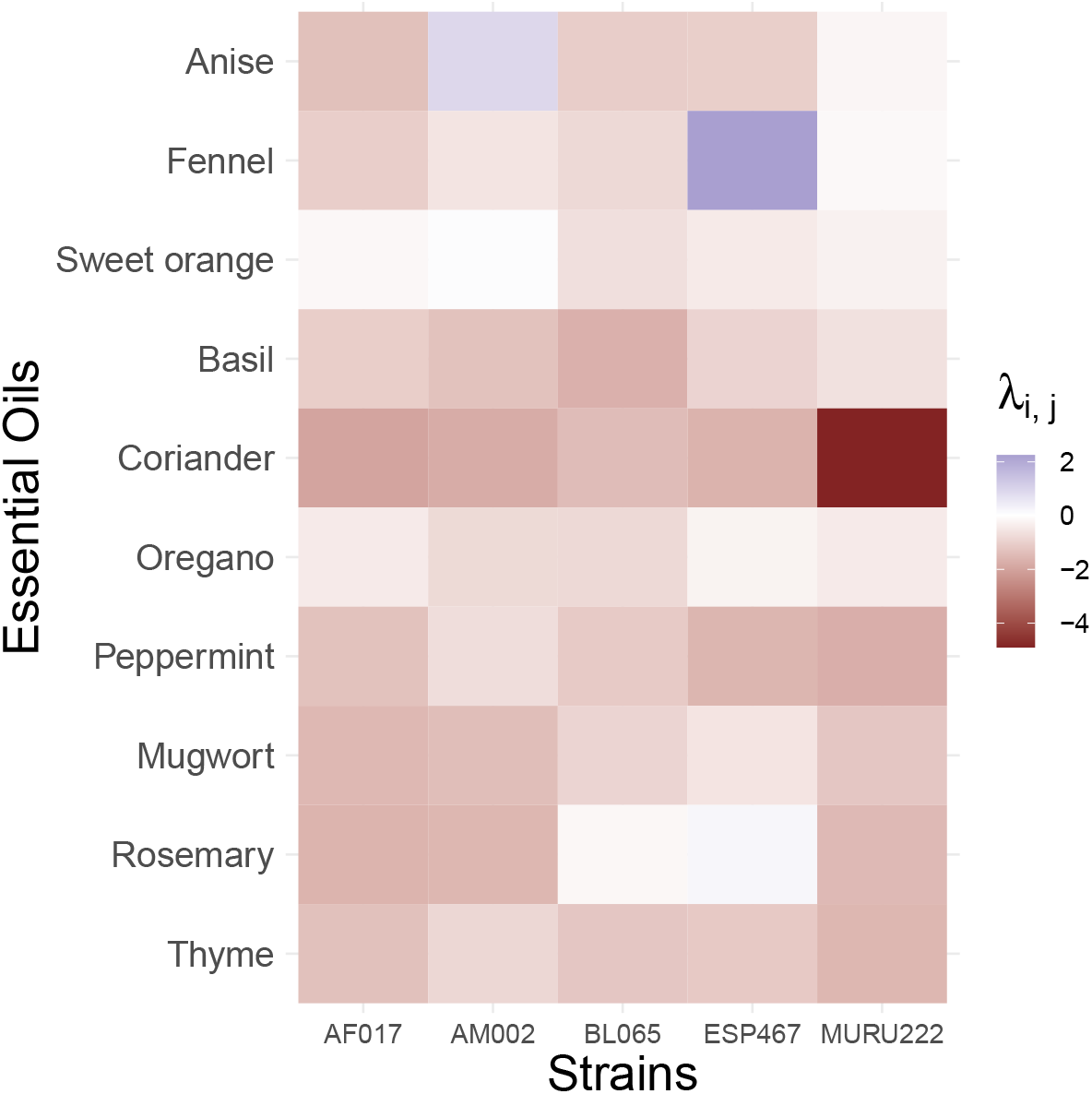
Effect of Essential Oils (rows) on the orientation of different strains of *T. evanescens* (columns). Parameter *λ*_*i,j*_ represent the effect of a given essential oil on the probability, for a parasitoid of a given strain, to stay in the same odor zone. Reddish and blueish squares respectively represent repulsion and attraction of a given strain for a given Essential Oil. The darker the square, the more intense is estimated the effect.

The presence of Basil, Coriander, Peppermint, Mugwort or Thyme EO tended to repel all strains (red squares in Fig 8). Indeed, negative values of *λ*_*i,j*_ mean that the probability to stay in a zone with odor was lowered by the presence of EO. As a result, the numbers of parasitoids in the zones containing odor drastically declined through time (Fig 9.a). For these five EOs, the posterior distribution of the average effect was clearly shifted to negative values (< 0.05% of positive values in Fig 10)

**Figure 9:**
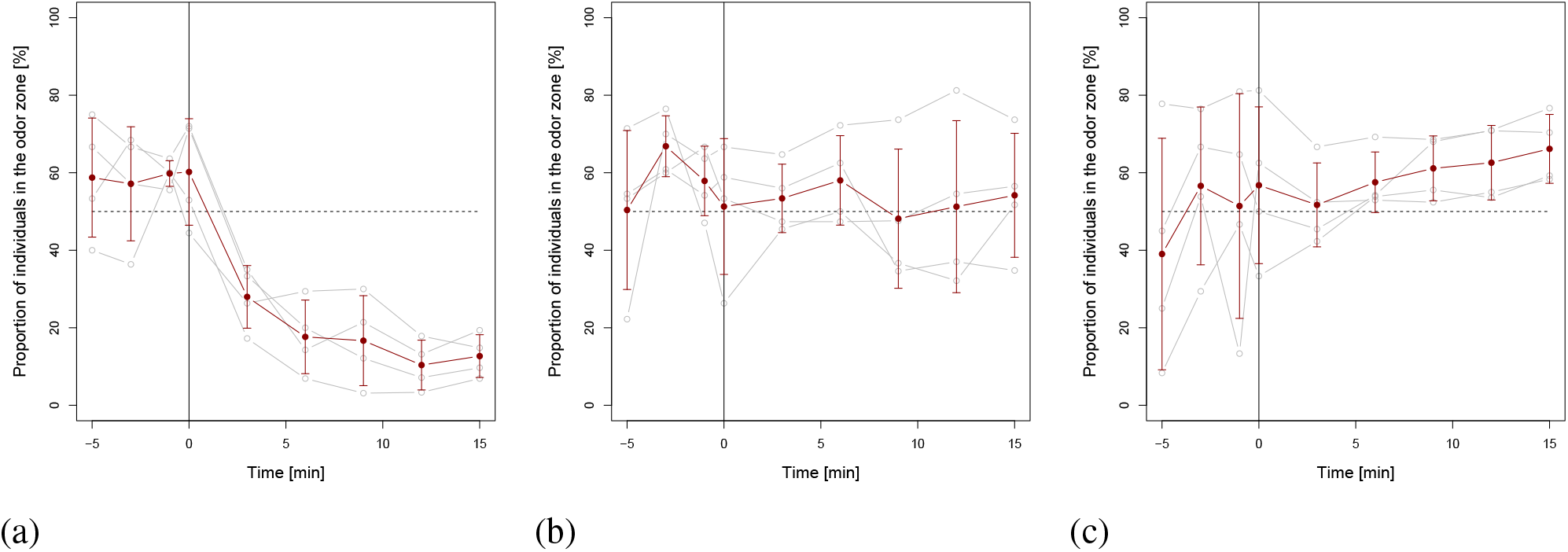
Proportion of individuals in the zone with odor during the experiment: (a) Coriander essential oil and MURU222 strain; (b) Sweet orange essential oil and AM002 strain; (c) Fennel essential oil and ESP467 strain. The black vertical bar represents the introduction of odor. Dashed line symbolize the random distribution (50% individuals). The four replicates are represented by the grey broken lines and summarized by their mean and standard deviation with the dark red broken lines and error bars.

**Figure 10:**
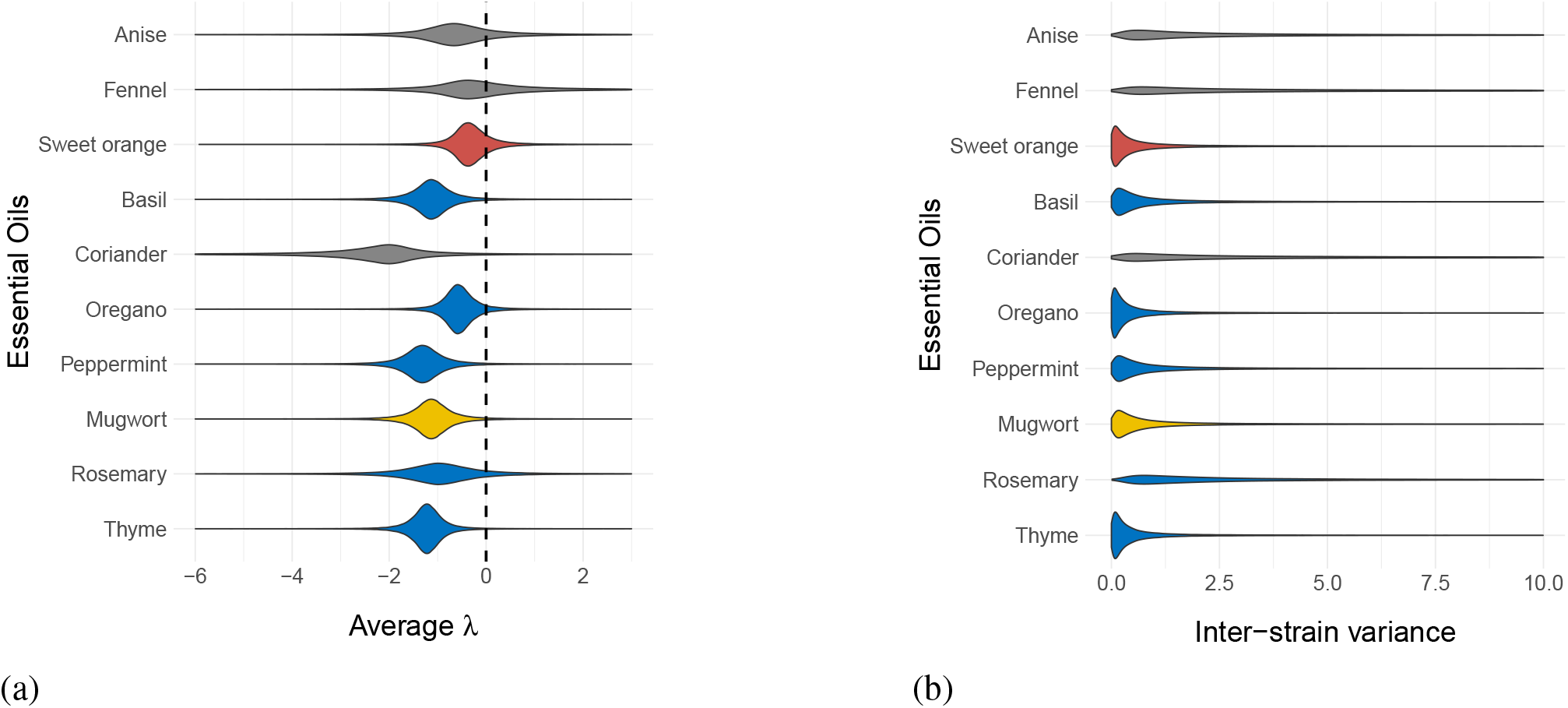
Estimated posterior distribution of the essential oil specific (a) mean (*m*_*λ*(*i*)_) and (b) inter-strain variance (*v*_*λ*(*i*)_) used to define parameter *λ*_*i,j*_ in model 1 from Table 4. Colors represent plant families: grey for Apiaceae, orange for Rutaceae, blue for Lamiaceae and yellow for Asteraceae. (a) Negative and positive values respectively represent global repulsiveness and attraction of a given Essential Oil. (b) Higher values mean larger differences in the essential oils effect between strains.

For Anise, Fennel, Sweet orange, Oregano or Rosemary, the impact on parasitoid behavior qualitatively differed between strains (Fig 8). On average, their presence decreased the probability for parasitoids to stay in a zone with odor. However, the posterior distribution of the average effect included 0 (Fig 10.a). Their presence could thus be almost neutral (white squares in Fig 8) and results in a nearly homogeneous distribution in the experimental setup (Fig 9.b). For both Anise and Fennel EOs, parameter *λ*_*i,j*_ were actually estimated as positive for at least one strain *j* (blue squares in Fig 8). This attractive effect resulted in an increasing proportion of individuals in the odor zones throughout the experiments (Fig 9.c).

Inter-strain variability was important. Three scenarios, defined by the shape of the posterior distribution of the EO-specific inter-strain variance (Fig 10.b), can be recognized: (i) For Sweet orange, Oregano and Thyme, inter-strain variance is very low (estimated mean of inter-strain variance < 1.6) and the effect of EOs on parasitoid behavior is very consistent for all tested strains. (ii) For Basil, Peppermint and Mugwort, results are slightly different between strains (estimated mean of inter-strain variance ∈ [1.6, 2.3]); and (iii) For Anise, Fennel, Coriander and Rosemary EOs, the variability of effect according to the tested strains is considerable (estimated mean of inter-strain variance > 6). There were different result across strains either quantitatively (for Coriander EO), or even qualitatively (for Anise, Fennel and Rosemary EOs).

## Discussion

We tested both toxicity and behavioral consequences of 10 essential oils (EOs) potentially used as biopesticides. These EOs all affected *T. evanescens* pre-imaginal survival, with some variability in the severity of their impact. Regarding the movement patterns of trichogramma adults, the majority of EOs had a repellent effect for naive females. In a few cases, however, the EOs seemed either neutral or even had a slight attractive effect. These results might be summarized by describing five groups of EOs (Table 5): (i) low effects: Rosemary and Sweet Orange EOs show low toxicity on pre-imaginal parasitoids and no observable impact on orientation behavior; intermediate effects: Mugwort essential oil is moderately toxic and seem repellent for naive females; (iii) contrasting effects: Oregano essential oil is highly toxic yet has almost no discernible effect on parasitoid behavior; (iv) variable effects: Anise and Fennel EOs are highly toxic for pre-imaginal parasitoid and can have repellent to attractive effect on females depending on strains; (v) strong effects: Basil, Coriander, Peppermint and Thyme EOs are highly toxic and invariably very repellent for *T. evanescens*. There was no correlation between the effect of EOs on parasitoid behavior and their toxicity on pre-imaginal survival (Fig 11; Pearson t(48)=0.63, p=0.53). We expected that the more toxic EOs might provoke greater avoidance and thus be more repellent (Araújo et al., 2020). Consistent with this expectation, EOs with low toxicity (Rosemary and Sweet Orange) showed little impact on parasitoid distribution. Similarly, some of the highly toxic EOs (Basil, Coriander, Peppermint and Thyme) were highly repellent. However, both Anise and Fennel EOs, that are highly toxic for pre-imaginal parasitoids, provoked no avoidance behavior, and for some strains, even appeared to be attractive (Fig 11).

**Table 5:**
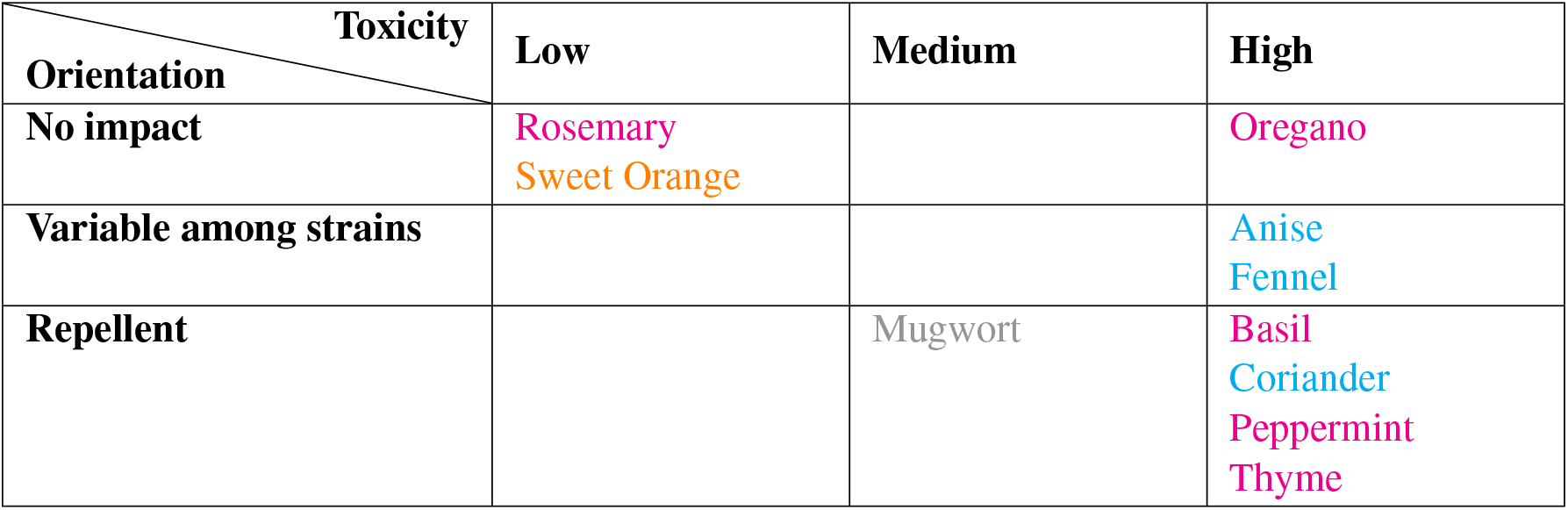
Summary of the results obtained regarding the toxicity of 10 EOs on *T. evanescens* pre-imaginal development and their effect on parasitoid orientation. Colors represent plant families: cyan for Apiaceae, grey for Asteraceae, magenta for Lamiaceae and orange for Rutaceae.

| **Toxicity** | **Low** | **Medium** | **High** |
| --- | --- | --- | --- |
| **Orientation** |  |  |  |
| **No impact** | Rosemary<br>Sweet Orange |  | Oregano |
| **Variable among strains** |  |  | Anise<br>Fennel |
| **Repellent** |  | Mugwort | Basil<br>Coriander<br>Peppermint<br>Thyme |

**Figure 11:**
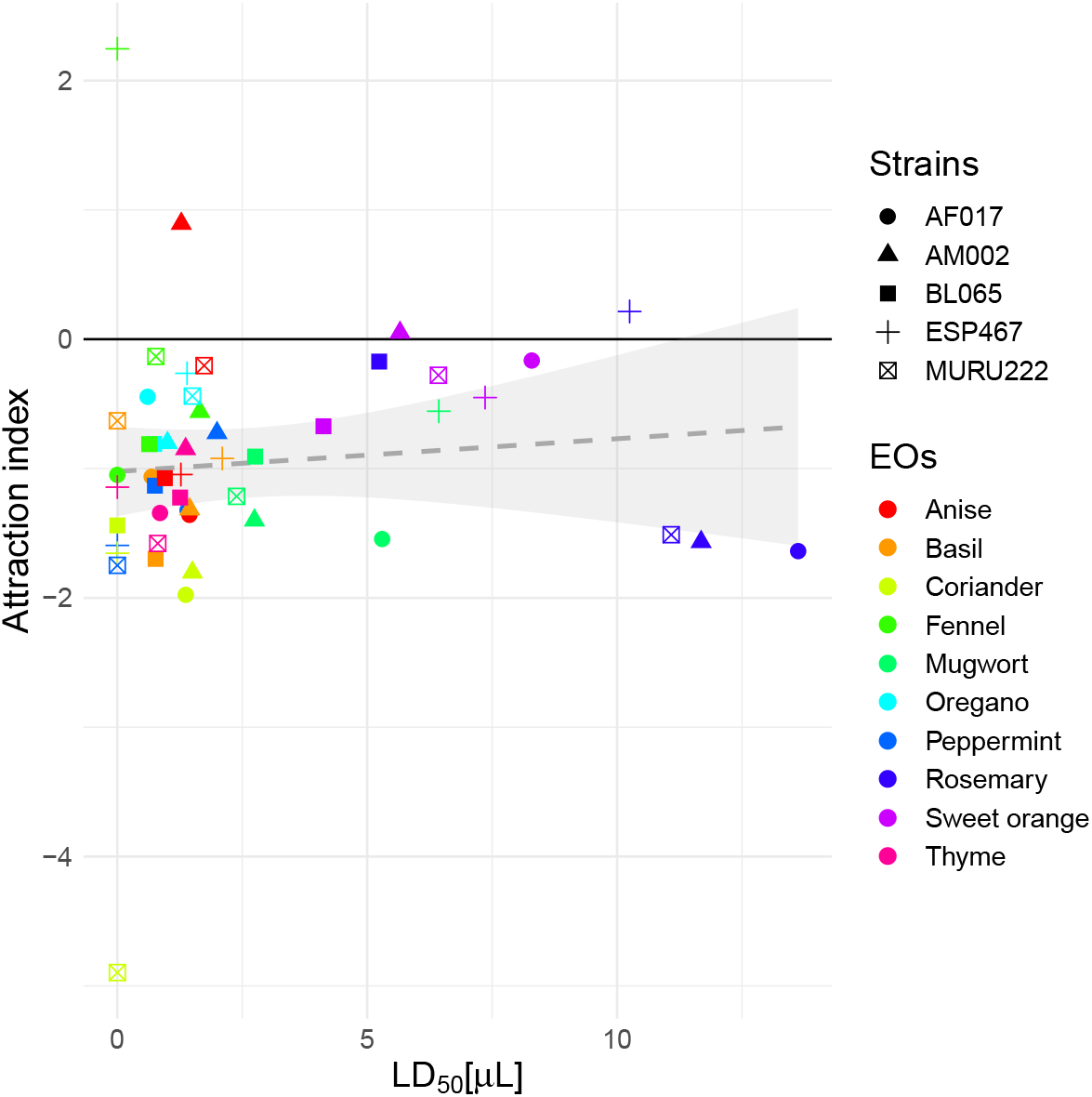
Lack of correlation between attraction and toxicity for the different essential oils (colors) and strains (symbols) tested. Attraction index is parameter *λ*_*i,j*_ estimates (see Fig 8). Toxicity on pre-imaginal developement is represented by the predicted LD_50_ (i.e. the dose at which 50% of the parasitized eggs do not emerge). Grey dashed line (and shadow) represent the estimation of a non-significant linear model (and standard error).

Pure EOs displayed fumigant toxicity on *T. evanescens* development. Indeed, all the tested EOs reduced the pre-imaginal survival of parasitized eggs. Anise, Fennel, Basil, Coriander, Oregano, Peppermint and Thyme appeared particularly toxic since they drastically reduced pre-imaginal development, even at low dose. The insecticidal potential of these EOs on non-target insects such as parasitoids is thus confirmed (Regnault-Roger, 1997; Sampson et al., 2005; Rani and Sandhyarani, 2012; Mossa, 2016). Some EOs are reported to act as Insect Growth Regulators and disrupt insect developement by inhibiting biosynthetic processes at different growth stages, thus reducing adults emergence (Agarwal et al., 2001; Kumar et al., 2011; Mossa, 2016). In *Trichogramma embryophagum* (Hartig) and *T. evanescens*, exposition of pre-imaginal stages to *Ferula assfoetida* essential oil increases pre-imaginal development time, decreases emergence rate and increases drastically wing abnormality among the emerged adults (Poorjavad et al., 2014). In our Bayesian analyses, we found that the impact on the second phase of development (pupal stage) was stronger than on the first phase (egg-larval stages). Our data better support this hypothesis than a potential accumulation effect of fumigant toxicity over time. It could confirm that pupal stages are more sensitive to exposition to EOs in *Trichogramma* (Parreira et al., 2018a,b). Indeed, the emergence of parasitized eggs of *E. kuehniella* was not affected by a five-second immersion in diluted EOs during the egg/larva stage (<1 day after parasitism) while it was reduced by more than 30% when immersion occurred at the pupal stage (7-8 days after parasitsm) for *T. pretiosum* (Parreira et al., 2018a) and *T. galloi* (Parreira et al., 2018b). Early instar parasitoids, that have very minute size, might be protected inside the host egg while after growth and host feeding (Volkoff et al., 1995), parasitoid pupae might be more directly exposed to air pollution.

Sweet Orange and Rosemary EOs were almost innocuous on pre-imaginal development. This difference is probably due to the mode of action of the main chemical compounds of these oils. Indeed, Rosemary EO is mainly composed of 1,8-cineole (or eucalyptol) (Suppl. Inf. S.1, see also (Isikber et al., 2006; Isman et al., 2008)). 1,8-cineole is known to inhibit the activity of acetylcholinesterase (AChE), an enzyme present in neuro-neuronal and neuro-muscular junctions (Mills et al., 2004; Jankowska et al., 2018). AChE inhibition causes paralisis and death of insects (Ryan and Byrne, 1988). At low dose, Rosemary essential oil may thus have little growth inhibition effet on *Trichogramma* as it was shown on beetles (Isikber et al., 2006) or noctuid caterpillars (Akhtar et al., 2008). Nevertheless, Rosemary EO is particularly toxic on adult stages (Isikber et al., 2006; Isman et al., 2008; Hanane et al., 2018). Moreover, both *Citrus aurantium* and *Rosemarinus officinaly* also repel phytophagous insects (Hori, 1998; Saeidi et al., 2011).

Most tested EOs affected the behavior of *T. evanescens* by increasing the probability to leave the odor zone of the olfactometer. Basil, Coriander, Peppermint, Mugwort or Thyme EOs seem particularly repellent for all tested strains. A repellent activity of EOs was recently documented in two *Trichogramma* species (Parreira et al., 2019; Alcántara-de la Cruz et al., 2021). In *T. pretiosum* (Parreira et al., 2019) and *T. galloi* (Alcántara-de la Cruz et al., 2021), previous exposition of host eggs to *Zingiber officinale, Allium sativum* and *Carapa guianensis* EOs, respectively, inhibited or drastically reduced parasitism rate. For both parasitoids, *Citrus sinensis, M. piperita, O. vulgare* or *T. vulgaris* EOs did not affect parasitism rate in this non-choice situation (Parreira et al., 2019; Alcántara-de la Cruz et al., 2021). In contrast, in our experimental setup, where individuals might choose either to be directly exposed to pure EOs or to escape and stay in an odorless zone, Peppermint and Thyme proved to be very repellent. Theses differences are probably explained by the mode of exposure to EOs, and by the dose used. In our experimental conditions, fumigant application of highly concentrated EOs is extremely repellent for *T. evanescens*. On the contrary, a past contact of host eggs to diluted solution of Peppermint and Thyme EOs seem not be so repulsive for parasitoids. Tested on the egg parasitoid *Trissolcus basalis*, three-day residues of pure *T. vulgaris* decreased parasitism rate while seven-day residues had no effect on parasitoid behavior (González et al., 2013). The effects of residues may decrease with both dose and time. Exposure mode is thus crucial to evaluate parasitoid susceptibility to EOs.

A crucial point using EOs is the dose of application. The relationship between the toxicity of EOs and the dose of application is commonly described with *doses* for ingestion, injection or contact bioassays and *concentrations* for inhalation toxicity. In our pre-imaginal survival experiment, we chose LD_50_ as dose descriptor since we only measured the substance dropped on the cotton plug. We used dose treatments (909, 1818 and 3636 ppm) closed to the estimated LC_01_, LC_10_, LC_50_ (877, 1758 and 4126 ppm) of 3 minutes exposition of *F. assafoetida* EO on adult *Trichogramma* wasps (Poorjavad et al., 2014). In our analyses, the predicted LD_50_ were far below these estimations since the corresponding LC_50_ would be lower than 365 ppm for the majority of the tested EOs, except Mugwort, Rosemary and Sweet orange EOs whose corresponding LC_50_ would remain below 2000 ppm. Long-lasting exposition of parasitized eggs to EOs might thus be more toxic than brief exposition of adult parasitoids. However, in our experiment, how the concentration of EOs inside the glass tube evolved over time remained unknown. The initial EO concentration could be affected in different ways. First, since tubes were not hermetically closed (cotton plugs), EOs might slightly evaporate. Second, with air contact and ambient temperature, EOs could oxidize and degrade some compounds (Turek and Stintzing, 2013). Third, some volatiles compounds from EOs could saturate the headspace inside the glass tubes (Rodrigues et al., 2003). This effect of saturation was probably observed in the olfactometry bioassays. Indeed, the lack of difference between zones exposed to air coming from the bottles with respectively 5*μL* and 10*μL* suggests that the odor airflow was probably very charged in EOs, at least from the parasitoid perspective. In a pest control context, the concentration of EOs are generally lower than the doses we used in our experiment. In aphid control for example, LC_50_ of efficient EOs are usually below 1 *μ*L.*L*^−1^ in fumigation applications and below 1 *μ*L.mL^−1^ in contact applications (Ikbal and Pavela, 2019). Our results might thus not directly predict what would happen in the field, but they highlight the need for EO evaluation in a tri-trophic context to test together the effect on the pest, the plant (Dunan et al., 2021) and the natural enemies.

The importance of inter-strain variability was particularly striking regarding orientation behavior (Milonas et al., 2009). Indeed, for Fennel, Anise, Sweet orange, Oregano and Rosemary EOs, the response of parasitoids was qualitatively different across strains. Some strains were repelled while other were indifferent, or even slightly attracted, by Anise and Fennel EOs. The main chemical compound of both Anise and Fennel EOs is anethole (Suppl. Inf. S.1). Anethole has biopesticidal potential (Sousa et al., 2021), and can be attractant for some insects such as scarabs (Toth et al., 2003) or lovebugs (Cherry, 1998). Anethole could thus act as an attractant for some strains of *T. evanescens*. However, strain AM002 was slightly attracted by Anise and slightly repelled by Fennel while ESP467 was attracted by Fennel and repelled by Anise. The effect of chemical composition on behavior must thus be more complex than the mere presence/absence of a single compound. Inter-strain variation might also result from the local adaptation of foraging behavior to different environmental conditions and host plants (Vos and Hemerik, 2003; Tamo et al., 2006). Strains AM002 and ESP467 were sampled respectively on quince and common bean. None of these plants (or fruits) contains anethole or other phenylpropanoid organic compounds (Tsuneya et al., 1983; Karolkowski et al., 2021). However, these plants might not be fully representative of the local environmental conditions of these strains. On the one hand, parasitoids were reared in the laboratory, without plants, for about 50 generations before experiments. On the other hand, the parasitoid sampling protocol, based on sentinel egg cards exposition, resulted in extremely low capture rate (<5%) and showed no reproducible patterns regarding host-plant associations (Ion Scotta, 2019). It might thus be difficult to connect parasitoid strains’ olfactory preferences with the volatile organic compounds of the sampled plants.

A noteworthy facet of our olfactory design is that parasitoid females were less than two days old, had probably mated (since they were reared with males). They had never been exposed to any olfactory stimulus except those from the substitute host from which they emerged (15-day old irradiated eggs of *E. kueniella*). Previous experiments on laboratory-reared *T. evanescens* showed that inexperienced females were not attracted to the synthetic sex pheromone of their hosts, contrary to females with previous oviposition experience (Schöller and Prozell, 2002). If some species of *Trichogramma* that were reared on factitious hosts were able to respond innately to native host cues (Milonas et al., 2009; Geetha, 2010), previous oviposition experience seems important for responding to olfactory signals (Kaiser et al., 1989; Fatouros et al., 2005; Consoli et al., 2010; Wilson and Woods, 2016). In this study, most EOs elicited escape behaviors in naive laboratory-reared females. This response might of course correspond to the avoidance of toxic compounds, but could also be induced by the perception of a strong unidentified olfactory stimulus, resulting in fear and avoidance due to neophobia (Corey, 1978). For generalist egg parasitoids such as *Trichogramma*, foraging decisions might greatly depend on learning abilities and on how infochemicals are linked to previous experience (Vet and Dicke, 1992; Wajnberg and Colazza, 2013; van Oudenhove et al., 2017).

This study confirms that fumigant application of EOs can have negative non-target effects on egg parasitoids *T. evanescens*, be it through mortality or repulsion. IPM programs must thus be extremely cautious in how and when to apply EOs if the program integrate biological control with parasitoids (González et al., 2013; Poorjavad et al., 2014). Indeed, EOs jeopardize the success of natural regulation by direct or indirect effects on natural enemy fitness. Using EOs for plant protection would require to carefully study the benefits (toxicity on pests) and risks (effects on natural enemies and phytotoxicity) (Dunan et al., 2021). This study also sets out possibilities for biocontrol programs that integrate semiochemicals to optimize the efficiency of egg parasitoids in agroecosystems (Wajnberg and Colazza, 2013). The potential of attraction of Anise and Fennel EOs on the one hand and the repulsion of *T. evanescens* for Basil, Coriander, Peppermint, Mugwort or Thyme EOs on the other hand might give clues for defining “push-pull” strategies based on the manipulation of natural enemies behavior with living plants or plant extracts (Khan et al., 2006).

## Acknowledgements

We thank the three PCI reviewers for their valuable comments. We thank Julien Papaix for his feedbacks on Bayesian analysis. We warmly thank Lydia Ottenwaelder for her help in some toxicity experiments. We are grateful to Robin Laugier for estimating *T. evanescens* sex-ratio. We thank Mélina Cointe and Nicolas Ris for helpful discussions and comments. The *Trichogramma* strains were provided by the Biological Resource Center EP-Coll that is a part of BRC4Env, the pillar “Environmental Resources” of the Research Infrastructure AgroBRC-RARe.

## Data, script and code availability

Data, R scripts and ImageJ macros are available in the INRAE data repository https://doi.org/10.15454/LSIPKT.

## Funding section

This work was financially supported by INRAE’s Department of Plant Health and Environment (project *Odor-amma*) and the Université Côte d’Azur (IDEX “Investissements d’Avenir UCAJEDI”, project reference ANR-15-IDEX-01).

