## Supplementary information for "Non-target effects of ten essential oils on the egg parasitoid *Trichogramma evanescens*"

### S.1 Essential Oils composition

The composition of the essential oils detailled in Table S.1 comes from the quality control performed by the supplier Esperis s.p.a. This composition can be graphically represented with a hierarchical cluster analysis using Ward's clusting criterion (Fig S.1) (Ward, 1963; Murtagh and Legendre, 2014).

Table S.1: Composition (in %) of the Essential Oil chemotypes according to the supplier Esperis s.p.a.. Bold values identify major compounds (>50%).

| Compounds \ Plants | Green anise | Fennel | Sweet orange | Basil | Coriander | Oregano | Peppermint | Mugwort | Rosemary | Thyme |
|---|---|---|---|---|---|---|---|---|---|---|
| alpha-pinene | 0 | [0.1, 0.5] | [0.1, 0.5] | 0 | [5, 7.5] | [1, 2.5] | [1, 3] | [1, 3] | [10, 12.5] | 0 |
| alpha-terpinene | 0 | 0 | 0 | 0 | 0 | [1, 2.5] | 0 | 0 | 0 | [2.5, 5] |
| alpha-terpineol | 0 | 0 | 0 | 0 | 0 | 0 | 0 | [1, 3] | 0 | 0 |
| alpha-thujone | 0 | 0 | 0 | 0 | 0 | 0 | 0 | **[60, 70]** | 0 | 0 |
| anethole | $\geq$ **90** | **[60, 70]** | 0 | 0 | 0 | 0 | 0 | 0 | 0 | 0 |
| beta-caryophyllene | 0 | 0 | 0 | 0 | 0 | 0 | 0 | [1, 3] | 0 | 0 |
| beta-myrcene | 0 | 0 | [1, 2.5] | 0 | 0 | [1, 2.5] | 0 | 0 | 0 | 0 |
| beta-pinene | 0 | 0 | 0 | 0 | 0 | 0 | 0 | 0 | [7.5, 10] | 0 |
| borneol | 0 | 0 | 0 | 0 | 0 | 0 | 0 | [1, 3] | 0 | 0 |
| camphene | 0 | 0 | 0 | 0 | 0 | 0 | 0 | 0 | [2.5, 5] | 0 |
| camphor | 0 | 0 | 0 | 0 | [3, 5] | 0 | 0 | [15, 20] | [5, 7.5] | 0 |
| carvacrol | 0 | 0 | 0 | 0 | 0 | **[70, 80]** | 0 | 0 | 0 | [10, 30] |
| cinnamaldehyde | 0 | 0 | 0 | 0 | 0 | 0 | 0 | [1, 3] | 0 | 0 |
| d-limonene | [0.1, 0.5] | [1, 3] | $\geq$ **90** | [0.5, 1] | [1, 3] | [0.5, 1] | [1, 3] | 0 | [5, 7.5] | [30, 40] |
| estragole | [1, 3] | [1, 3] | 0 | **[70, 80]** | 0 | 0 | 0 | 0 | 0 | 0 |
| eucalyptol | 0 | 0 | 0 | [1, 3] | 0 | 0 | [3, 5] | [7.5, 10] | [40, 50] | 0 |
| eugenol | 0 | 0 | 0 | [0.5, 1] | 0 | 0 | 0 | [1, 3] | 0 | 0 |
| geraniol | 0 | 0 | 0 | 0 | [1, 3] | 0 | 0 | [1, 3] | 0 | 0 |
| geranyl acetate | 0 | 0 | 0 | 0 | [1, 3] | 0 | 0 | 0 | 0 | 0 |
| linalool | 0 | 0 | [0.1, 0.5] | [7.5, 10] | **[60, 70]** | 0 | 0 | 0 | [1, 2.5] | [1, 2.5] |
| menthol | 0 | 0 | 0 | 0 | 0 | 0 | **[50, 60]** | 0 | 0 | 0 |
| menthone | 0 | 0 | 0 | 0 | 0 | 0 | [20, 30] | 0 | 0 | 0 |
| menthyl acetate | 0 | 0 | 0 | 0 | 0 | 0 | [5, 7.5] | 0 | 0 | 0 |
| methyl eugenol | 0 | 0 | 0 | [0.5, 1] | 0 | 0 | 0 | 0 | 0 | 0 |
| p-cymene | 0 | 0 | 0 | 0 | [1, 3] | [7.5, 10] | 0 | 0 | 0 | [10, 15] |
| pulegone | 0 | 0 | 0 | 0 | 0 | 0 | [1, 3] | 0 | 0 | 0 |
| terpinen-4-ol | 0 | 0 | 0 | 0 | 0 | [2.5, 5] | 0 | 0 | 0 | 0 |
| terpineol | 0 | 0 | 0 | 0 | 0 | 0 | 0 | 0 | [1, 2.5] | 0 |
| thymol | 0 | 0 | 0 | 0 | 0 | [1, 2.5] | 0 | 0 | 0 | [30, 40] |

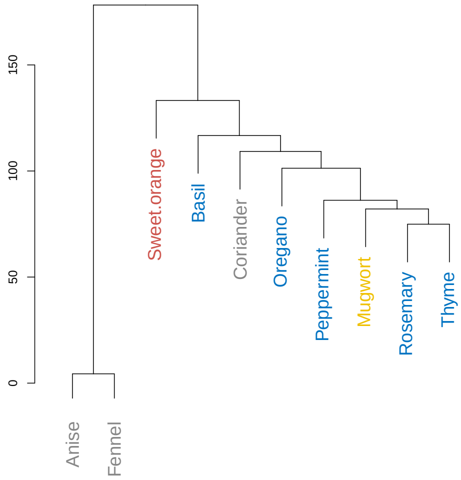


Figure S.1: Hierarchical cluster on Essential Oils composition using Ward's criterion. Chemical compositions are documented with confidence intervals by the supplier (see Table S.1). The values used for this graphical representation are the mean of the intervals normalized to 100%. Colors represent plant families: grey for Apiaceae, orange for Rutaceae, blue for Lamiaceae and yellow for Asteraceae.

### S.2 Simulation of air flow in the olfactometer

To determine the nature of the airflow within the olfactometer, we used numerical simulation using Computational Fluid Dynamics (CFD) model. This modelling provides information on the internal air velocity fields based on the numerical resolution of Navier-Stokes equations describing the air flow.

The air in the olfactometer is assimilated to an incompressible fluid governed by Navier-Stokes equation:

$$\frac{\partial(U\Phi)}{\partial X} + \frac{\partial(V\Phi)}{\partial Y} + \frac{\partial(W\Phi)}{\partial Z} = \Gamma.\bigtriangledown^2 \Phi + S_\Phi$$

where $\Phi$ represents the three-dimensional momentum (Navier-Stokes); $U$, $V$ and $W$ are the components of the velocity vector; $\Gamma$ is the diffusion coefficient; and $S_\Phi$ is the source term.

Advanced CFD software (Ansys Fluent ©) was used to solve this highly non-linear equation using a spatial finit volume discretization. The olfactometer's computational domain is discretized according to a Cartesian body fitted coordinates with refinement close to all solid walls. Velocity inlet boundary conditions were used to define the flow velocity, along with all relevant scalar properties of the flow. The boundary conditions at the outlet are automatically calculated to satisfy the continuity equation. The visualization interface of the CFD software allows to display the dynamic fields distributed within the olfactometer (Fig S.2).

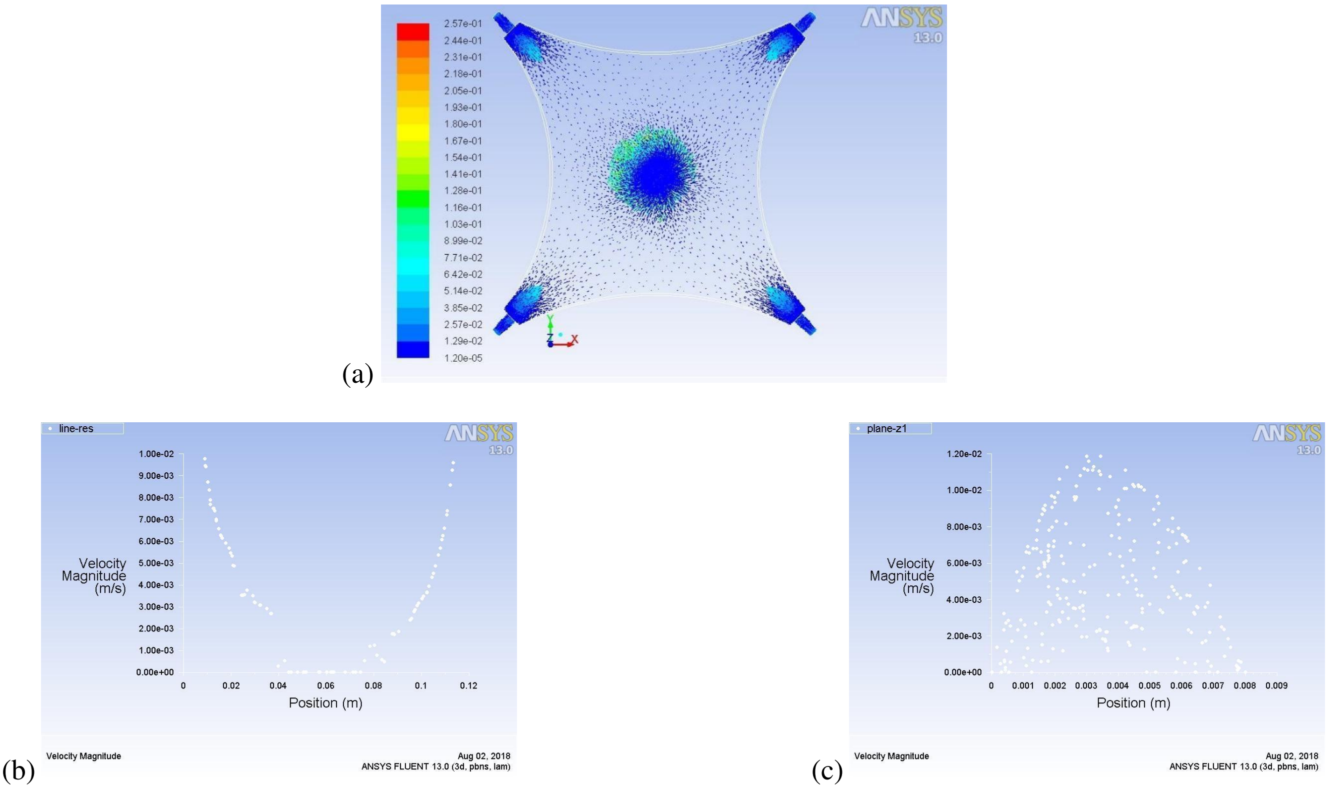


Figure S.2: Airflow modeling inside the exposure chamber of the four-arms olfactometer. (a) Overview of the air velocity; (b) Air velocity from the inlet to the central hole; (c) Air velocity in the olfactometer thickness.

The modelling results, as shown in Fig S.2.a, indicate that the air flow at the central hole is laminar without turbulence inside the tube. At the extremity of the four-arms of the olfactometer, the air velocity increases due to the widening of the walls. At the outlet, an acceleration of the air velocity is observed due to the reunification of the four flows through the small diameter of the outlet tube (Venturi effect) (Fig S.2.a and S.2.b). With a flow rate of $1.6L.h^{-1}$ at each inlet, the air flow is uniformly distributed in the exposure chamber and no turbulence is observed. Regarding the air velocity through the thickness of the olfactometer ($0.8cm$) (Fig S.2.c), the air velocity at the walls is zero and increases in the center.

### S.3 Pre-imaginal survival

#### S.3.1 Estimation of the number of eggs

In step1-model (see Fig S.3 for model structure), all parameters were considered as unknown random variables with non-informative prior probability distributions: $\gamma_{B_0} \sim \mathcal{N}(0, 10^4), \gamma_{W_0} \sim \mathcal{N}(0, 10^4), \theta_{B_0} \sim \mathcal{N}(0.01, 10^4)$, $\theta_{W_0} \sim \mathcal{N}(0.01, 10^4), \sigma_{\gamma,B_0} \sim \mathcal{U}(0, 100), \sigma_{\gamma,W_0} \sim \mathcal{U}(0, 100), \sigma_{\theta,B_0} \sim \mathcal{U}(0, 100)$ and $\sigma_{\theta,W_0} \sim \mathcal{U}(0, 100)$. Three independent Markov Chain Monte Carlo (MCMC) chains were run in parallel. After an initial burn-in period of 5,000 iterations, the Bayesian algorithm was run 15,000 iterations and the corresponding sample of parameter posterior distributions were recorded. According to the Gelman-Rubin diagnostic (Gelman and Rubin, 1992), this model reached convergence ($\hat{R} < 1.01$).

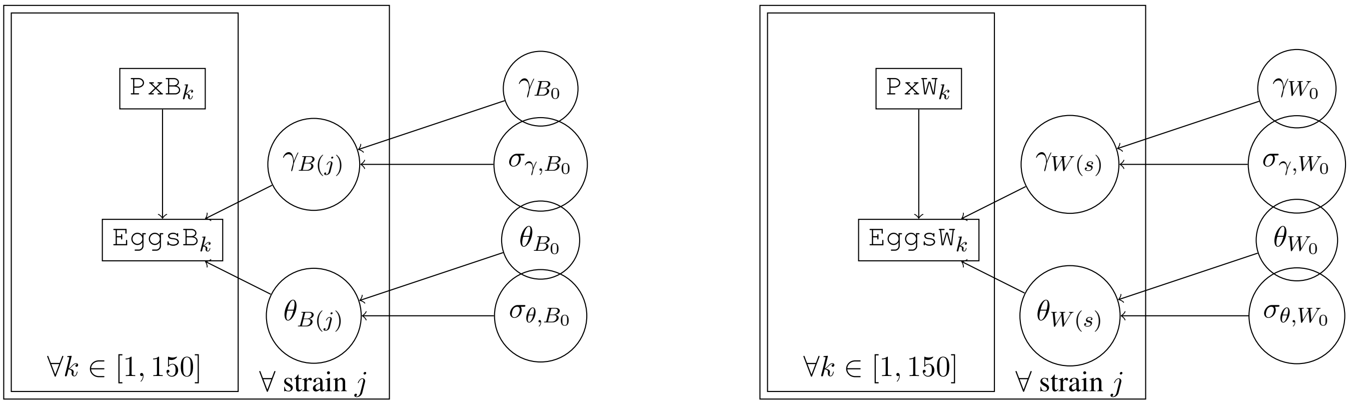


Figure S.3: Directed Acyclic Graph of the step1-model for either black (left pannel) or white (right pannel) eggs. For each replicate $k \in [1, 150]$, the number of eggs depends on the number of pixels of the corresponding color, such as $\texttt{EggsB}_k \sim \mathcal{P}(\gamma_{B(j_k)} + \theta_{B(j_k)}\texttt{PxB}_k)$ and $\texttt{EggsW}_k \sim \mathcal{P}(\gamma_{W(j_k)} + \theta_{W(j_k)}\texttt{PxW}_k)$. Small variation according to strains $j$ are described by the following links: $\gamma_{B(j)} \sim \mathcal{N}(\gamma_{B_0}, \sigma_{\gamma,B_0})$, $\gamma_{W(j)} \sim \mathcal{N}(\gamma_{W_0}, \sigma_{\gamma,W_0})$, $\theta_{B(j)} \sim \mathcal{N}(\theta_{B_0}, \sigma_{\theta,B_0})$ and $\theta_{W(j)} \sim \mathcal{N}(\theta_{W_0}, \sigma_{\theta,W_0})$. Arrows stand for probabilistic links. Circle nodes are unknown parameters and rectangle nodes are observed.

These parameter estimations allowed the number of eggs, either black or white, to be accurately predicted from the number of pixels of the respective color (Fig S.4). The number of black and white eggs, $EggsB$ and $EggsW$, were thus predicted from the number of pixels for all the 310 patches and round to unit. In a few cases (about 3%), the estimated number of black eggs being smaller than the number of emerging adults, the maximum between these two values was preferred in order to avoid convergence issues in the survival model.

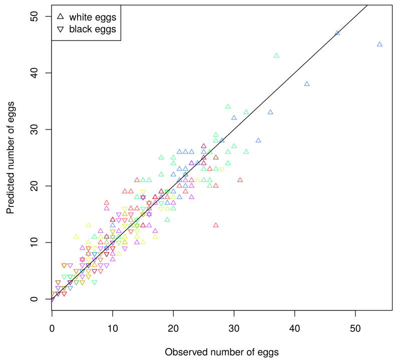


Figure S.4: Correlation between the number of eggs predicted by the number of pixels in a patch and the number of eggs observed. Spearman correlation values were 0.96 for black eggs and 0.91 for white eggs. Triangle shape codes either for black and white eggs. Colors represent the different strains. Black line is the strict equivalence (linear equation $y = x$).

#### S.3.2 Survival model

In the survival-model (see Fig S.5 for model structure), we used large non-informative distributions: for each strain $j$, mean parasitism rate $\kappa_j \sim \mathcal{B}eta(1,1)$, natural mortality $\delta \sim \mathcal{B}eta(1,1)$ and priors for sensitivity to essential oils are $\alpha_{i,j} \sim \mathcal{N}(0,10^4)$ and $\beta_{i,j} \sim \mathcal{N}(0,10^4)$ for each strain $j$ and essential oil $i$. To avoid survival probability to be higher than 1, parameters $\alpha_{i,j}$ and $\beta_{i,j}$ were constrained such as $\alpha_{i,j} > 0$ and $\beta_{i,j} > -\alpha_{i,j}$.

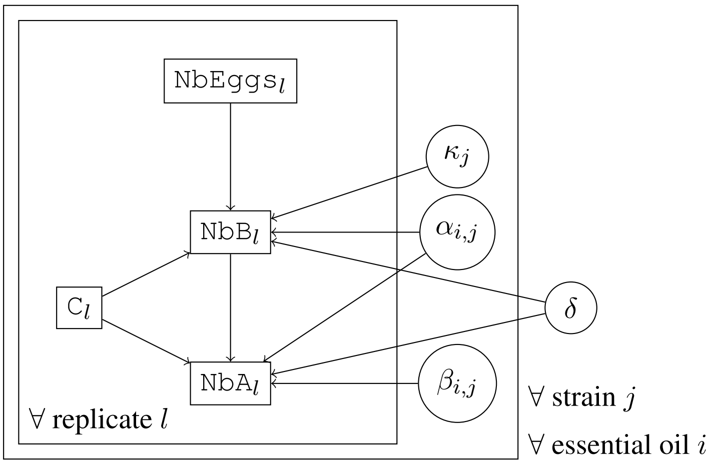


Figure S.5: Directed Acyclic Graph of the survival-model representing, for each patch $l$, the links between the number of eggs ($\texttt{NbEggs}_l$), the number of black eggs on fifth day ($\texttt{NbB}_l \sim \mathcal{B}in(\texttt{NbP}_l, \kappa_j(\delta e^{-\alpha_{i,j}C_l})^5)$) and the number of emerging adults ($\texttt{NbA}_l \sim \mathcal{B}in(\texttt{NbB}_l, (\delta e^{-(\alpha_{i,j}+\beta_{i,j})C_l})^9)$). $\texttt{C}_l$ is the essential oil dose at which patch $l$ was exposed. Parasitism rate depends on a mean parasitism rate of a given strain $\kappa_j$. $\delta$ is the natural survival rate. $\alpha_{i,j}$ and $\beta_{i,j}$ represent the effect of essential oils on survival. Simple or double arrows stand respectively for probabilistic or deterministic links. Circle nodes are unknown parameters and rectangle nodes are observed data.

Three independent MCMC chains were run in parallel for each model. After an initial burn-in period of 5,000 iterations, the Bayesian algorithm was run 45,000 iterations and the corresponding sample of parameter posterior distributions were recorded. Chain convergence was studied for the different tested models. All parameters reached convergence according to Gelman-Rubin diagnostic ($\hat{R} < 1.01$, Gelman and Rubin (1992)).

### S.4 Olfactometry bioassays

The olfactometry bioassays model was built as represented in the directed acyclic graph (Fig S.6).

The parasitoids' repartition model was first fitted on data from control experiments (without odor). Parameters were considered as unknown random variables with non-informative prior probability distributions: $\mu \sim \mathcal{B}eta(1,2)$ ($\mu \in [0,1]$). In the different models (Table 3), different parameter $\psi$ were tested such as: independent $\psi_{-\infty} : S_z(t) = 0 \forall$ time $t, \forall$ zone $z$, constant $\psi \sim \mathcal{N}(0,10^4)$, strain-specific $\psi_j \sim \mathcal{N}(0,10^4) \forall$ strain $j$, experiment-specific $\psi_k \sim \mathcal{N}(m_\psi, v_\psi) \forall$ experiment $k (k \in [1,200])$ with $m_\psi \sim \mathcal{N}(0,10^4)$ and $v_\psi \sim \mathcal{U}(0,100)$, experiment-specific with strain-variability $\psi_{k(j)} \sim \mathcal{N}(m_{\psi(j)}, v_\psi) \forall$ experiment $k(k \in [1,200])$ with $m_{\psi(j)} \sim \mathcal{N}(0,10^4) \forall$ strain $j$ and $v_\psi \sim \mathcal{U}(0,100)$. Three independent Markov Chain Monte Carlo (MCMC) chains were run in parallel. After an initial burn-in period of 5,000 iterations, the Bayesian algorithm was run 45,000 iterations and the corresponding sample of parameter posterior distributions were recorded. According to the Gelman-Rubin diagnostic (Gelman and Rubin, 1992), this model reached convergence ($\hat{R} < 1.01$).

The parameter estimation of the best model selected in the control experiment analysis was used as prior to analyze data from treatment experiment (with odor). $\mu$ and $psi$ were thus modeled using the mean and the variance estimated in the previous step. Parameters $\omega$ and $\lambda$ were considered as unknown random variables

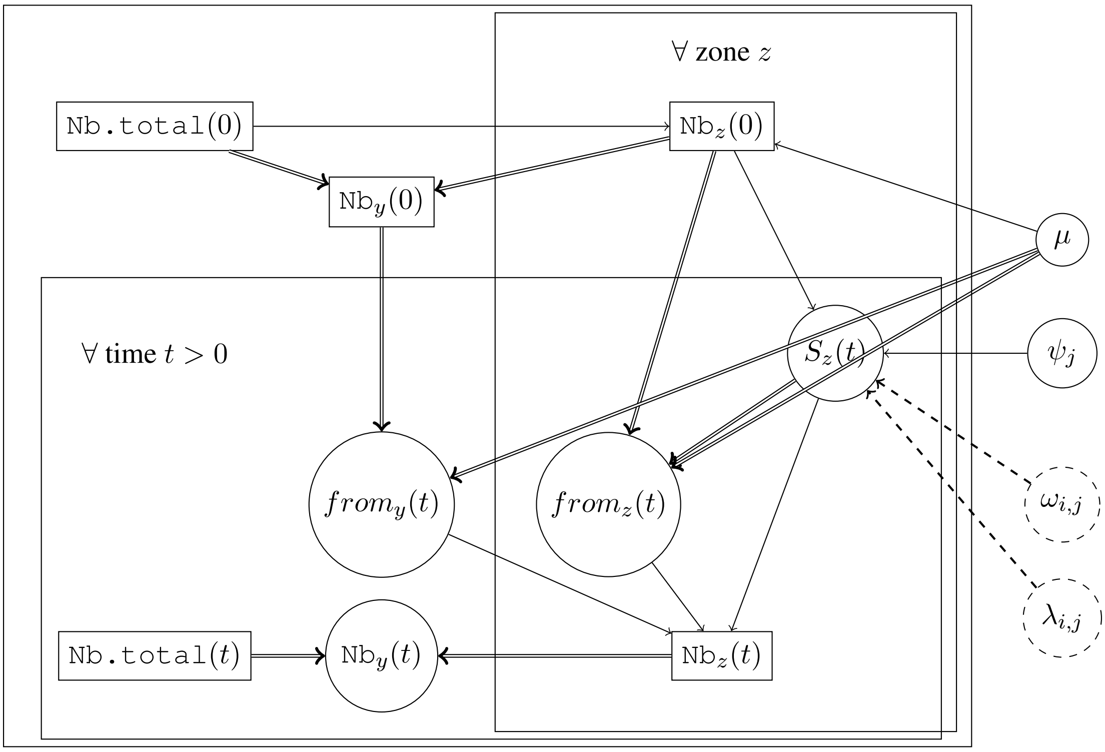


Figure S.6: Directed Acyclic Graph of the olfactometry bioassay model representing the links between the observed data (rectangle nodes) and the estimated variables and parameters (circles). For time $t = 0$ and other time $t$ ($t \in \{0, 2, 4\}$ for control experiments and $t \in \{0, 3, 6, 9, 12, 15\}$ for treatment experiments), $\texttt{Nb.total}(t)$ is the number of individuals in the exposure chamber; $\texttt{Nb}_z(t)$ and $\texttt{Nb}_y(t)$ are respectively the number of parasitoids in a given zone $z$ ($z \in \{1, 2, 3, 4\}$) or elsewhere in the exposure chamber; $S_z(t)$ is the number of individuals staying in a zone $z$ between $t$ and $t + 1$; and $from_z(t)$ is the flow of individuals arriving from a zone $z$ in another zone. $\mu$ is the probability that an individual would not be in a defined zone $z$ of the exposure chamber assuming a random distribution (see Fig 3). $\psi_j$ allows to define the probability for an individual to stay in a given zone (without odor) during $m$ min $\frac{1}{(1+e^{-\psi_j})^m}$. Parameter $\omega_{i,j}$ and $\lambda_{i,j}$ respectively represent the effect of 10 and 5 mL of essential oil $i$ on strain $j$: in odor zone, the probability of an individual to stay during $m$ min are respectively $\frac{1}{(1+e^{-(\psi_j+\omega_{i,j})})^m}$ and $\frac{1}{(1+e^{-(\psi_j+\lambda_{i,j})})^m}$ for 10 and 5mL of essential oil. $i$ and $j$ respectively being the essential oil and the strain of a given experiment. Simple or double arrows stand respectively for probabilistic or deterministic links. Dashed links represent links that are only taken into account for odor zones. Equation details are given in section .

with non-informative prior probability distribution. Different parameters were tested such as: $\lambda_0 = \omega_0 = 0$; $\lambda_i = \omega_i \sim \mathcal{N}(0, 10^4)$, $\forall i$; $\lambda_{i,\sigma(j)} = \omega_{i,\sigma(j)} \sim \mathcal{N}(m_{\lambda(i)}, v_\lambda)$ with $m_{\lambda(i)} \sim \mathcal{N}(0, 10^4)$ $\forall i$ and $v_\lambda \sim \mathcal{U}(0, 100)$; $\lambda_{i,\sigma_i(j)} = \omega_{i,\sigma_i(j)} \sim \mathcal{N}(m_{\lambda(i)}, v_{\lambda(i)})$ with $m_{\lambda(i)} \sim \mathcal{N}(0, 10^4)$ and $v_{\lambda(i)} \sim \mathcal{U}(0, 100)$ $\forall i$; $\lambda_{i,\sigma_i(j)} = \frac{1}{2} \cdot \omega_{i,\sigma_i(j)} \sim \mathcal{N}(m_{\lambda(i)}, v_{\lambda(i)})$ with $m_{\lambda(i)} \sim \mathcal{N}(0, 10^4)$ and $v_{\lambda(i)} \sim \mathcal{U}(0, 100)$ $\forall i$; $\lambda_{i,\sigma_i(j)} \sim \mathcal{N}(m_{\lambda(i)}, v_{\lambda(i)})$ with $m_{\lambda(i)} \sim \mathcal{N}(0, 10^4)$ and $v_{\lambda(i)} \sim \mathcal{U}(0, 100)$ $\forall i$ and $\omega_{i,\sigma_i(j)} \sim \mathcal{N}(m_{\omega(i)}, v_{\omega(i)})$ with $m_{\omega(i)} \sim \mathcal{N}(0, 10^4)$ and $v_{\omega(i)} \sim \mathcal{U}(0, 100)$ $\forall i$.

For each tested model, three independent MCMC chains were run in parallel for 50,000 iterations (with an initial burn-in period of 5,000 iterations). The corresponding sample of parameter posterior distributions were recorded. According to the Gelman-Rubin diagnostic (Gelman and Rubin, 1992), all estimated parameters in the tested models reached convergence ($\hat{R} < 1.01$).
